# Mitochondrial proteostasis mediated by CRL5^Ozz^ and Alix maintains skeletal muscle function

**DOI:** 10.1101/2023.07.11.548601

**Authors:** Yvan Campos, Ricardo Rodriguez-Enriquez, Gustavo Palacios, Diantha Van de Vlekkert, Xiaohui Qiu, Jayce Weesner, Elida Gomero, Jeroen Demmers, Tulio Bertorini, Joseph T. Opferman, Gerard C. Grosveld, Alessandra d’Azzo

## Abstract

High energy-demanding tissues, such as skeletal muscle, require mitochondrial proteostasis to function properly. Two quality-control mechanisms, the ubiquitin proteasome system (UPS) and the release of mitochondria-derived vesicles, safeguard mitochondrial proteostasis. However, whether these processes interact is unknown. Here we show that the E3 ligase CRL5^Ozz^, a member of the UPS, and its substrate Alix control the mitochondrial concentration of Slc25A4, a solute carrier that is essential for ATP production. The mitochondria in *Ozz*^−/−^ or *Alix^−/−^*skeletal muscle share overt morphologic alterations (they are supernumerary, swollen, and dysmorphic) and have abnormal metabolomic profiles. We found that CRL5^Ozz^ ubiquitinates Slc25A4 and promotes its proteasomal degradation, while Alix facilitates SLC25A4 loading into exosomes destined for lysosomal destruction. The loss of Ozz or Alix offsets steady-state levels of Slc25A4, which disturbs mitochondrial metabolism and alters muscle fiber composition. These findings reveal hitherto unknown regulatory functions of Ozz and Alix in mitochondrial proteostasis.

## INTRODUCTION

Skeletal muscle, one of the most specialized tissues in mammals, is made up of multinucleated cells called myofibers that are programmed to contract. The intricate architecture and dynamics of muscle fibers rely on their myosin composition and metabolic activity^1, 2^. During muscle contraction, the energy demand of the tissue is supplied by mitochondria that convert nutrients into ATP through the process of oxidative phosphorylation (OXPHOS). In skeletal muscle, mitochondria are categorized as subsarcolemmal or intermyofibrillar based on their position within the fiber, and they form an interconnected reticular membrane network that has been compared to the grid of a power plant^3, 4^. Subsarcolemmal mitochondria exhibit remarkable plasticity to adapt their volume; in contrast, intermyofibrillar mitochondria are less plastic but have higher rates of protein synthesis, enzyme activities, and respiration^3, 4^. For optimal cellular performance, these organelles must maintain a fine-tuned equilibrium between protein synthesis and degradation.

Mitochondrial proteostasis depends on the coordinated activity of several quality control (QC) mechanisms that regulate protein degradation to ensure a functional mitochondrial proteome. In mitochondria, only about 13 of the approximate 1500 proteins are encoded by mitochondrial DNA^5^; the remainder are synthesized in the cytosol and chaperoned into the mitochondria, where they are further distributed to sub compartments, mostly by two multiprotein complexes: Tom (translocase of the outer membrane) and Tim (translocase of the inner membrane)^3, 6–8^. In response to physiologic or pathologic stimuli, specific QC mechanisms come into play. These include mitochondrial proteases that are present at both the cytosolic and mitochondrial sites, the ubiquitin proteasome system (UPS), and the release of cargo-loaded vesicles, such as mitochondrial-derived vesicles (MDVs) or exosomes, destined for endolysosomal catabolism^9^. If any of these systems do not work properly, misfolded, damaged, or aggregated mitochondrial proteins accumulate, resulting in cellular dysfunction and disease, including several mitochondrial myopathies and cardiomyopathies^10–12^.

The UPS mediates the turnover of approximately 62% of the total mitochondrial proteome^13^, which includes resident proteins (49%) and proteins that are present in other organelles^14^. Chief examples of mitochondrial proteins that are targeted to the UPS for degradation are mitochondrial precursors, proteins responsible for OXPHOS^15, 16^, and those that maintain the morphology of the organelles (e.g., Fis1, Drp1, Mfn1, Mfn2, and MiD49). Several ubiquitinated mitochondrial proteins, for which the dedicated ubiquitin ligase has been identified, reside in the outer mitochondrial membrane (OMM) and are strategically placed close to the proteasome^13, 17^. For many others, especially those residing in the inner mitochondrial membrane^18^ and the matrix, the responsible ubiquitin ligases and mechanisms by which they are made accessible to the proteasomal machinery remain unknown^16^.

The UPS is selective; it targets only one ubiquitinated protein at a time. In contrast, MDVs and exosomes are loaded with mitochondrial proteins of different derivations, most of which are destined for endolysosomal degradation, albeit the mechanisms that coordinate their sorting into these nanovesicles have not been elucidated. Among the MDV cargoes are components of the TOM complex and subunits of the mitochondrial electron transport chain^19^. Members of the TOM complex, mitochondrial solute carriers (Slc), and other proteins of mitochondrial origin also have been identified in exosomes^20–23^. Thus, whether and to what extent MDVs resemble exosomes, in terms of their origin, characteristics, and function, remains to be determined^22^.

Beside these QC systems, cells utilize mitophagy as a surveillance mechanism to dispose of irreversibly damaged or dysfunctional mitochondria in bulk^24–27^. This regulated process involves UPS activation, which is canonically initiated by posttranslational modifications and association with PINK1 and Parkin^24–27^. PINK1 and Parkin also have been implicated in the biogenesis of MDVs, a finding that may connect the formation of MDVs and their endolysosomal degradation with mitophagy^22, 28, 29^. Thus, different mitochondrial QC mechanisms are most likely interconnected, but how they interact remains unexplored, especially in skeletal muscle.

In previous studies, we identified Ozz as the substrate-recognition component of CRL5^Ozz^, a muscle-specific Cullin RING–type ubiquitin ligase complex comprising Elongin B/C, Rbx1, and Cul5^30, 31^. CRL5^Ozz^ plays a role in myofiber differentiation and maturation during embryogenesis and muscle regeneration. Ozz, also known as Neurl2 (Neuralized-like protein 2), belongs to the SOCS (suppressor of cytokine signaling) family of proteins. The primary structure of Ozz includes two NHR (neuralized homologous repeat) domains that recognize substrate, and a SOCS box at the C terminus that interacts with Elongin B/C^31^. During early embryogenesis, Ozz is detected in the somites, at the tips of myotomal cells, near the intermyotomal septum, and in the myotendinous junctions^31^. During myofiber differentiation, Ozz expression increases progressively: it localizes initially to the tips of the differentiating myocytes and later throughout the mature fiber^31^. This expression pattern reflects the location of CRL5^Ozz^ substrates, which include canonical sarcomeric proteins, membrane proteins, and the ubiquitously expressed protein Alix or PDCD6IP^32^.

Alix is a multidomain scaffold protein that is a component of two large multiprotein complexes, the ESCRT (endosomal sorting complexes required for transport) and the F-actin cytoskeleton^32–42^. In this capacity, Alix directly or indirectly coordinates a myriad of cellular processes, including endocytosis, membrane repair/remodeling, viral growth, cytokinesis, multivesicular body sorting, and exosome biogenesis^32, 33, 35–38, 40, 42, 43^.

We have reported earlier that silencing *Alix* expression in C2C12 myoblasts alters the levels and distribution of F-actin, reduces the formation of membrane protrusions, and the release of extracellular vesicles^32, 44^. The interplay between these roles of Alix became apparent during the characterization of the *Alix^−/−^* mouse model^34^. These mice develop a severe brain phenotype, i.e., bilateral hydrocephalus, that we have linked to the improper assembly and positioning of an actomyosin-tight junction complex of the choroid plexus epithelial cells. This complex defines a spatial membrane domain that is essential for the maintenance of epithelial cell polarity and the epithelial barrier^34^. In this process, Alix serves as the bridge between membrane multiprotein complexes and the F-actin cytoskeleton^34^. However, the potential consequences of Alix deficiency in skeletal muscle have not yet been determined.

Here we explored whether Ozz and Alix mutually maintain the structural and functional integrity of skeletal muscle. Using genetic, proteomic, and imaging methods, we discovered that Ozz and Alix localize and function in mitochondria. By regulating the degradation of the solute carrier SLC25A4 (also known as ANT1), they cooperatively control mitochondrial proteostasis in skeletal muscle.

## RESULTS

### Ozz and Alix regulate mitochondrial morphology

To begin assessing whether ablating Alix in skeletal muscle alters the tissue’s phenotype and whether those muscles resembled the skeletal muscle in *Ozz^−/−^* mice^31^, we first examined the expression levels and spatial distribution of Alix, compared to Ozz, in lysates from different skeletal muscle types (Figure S1A). Although we detected no obvious differences in the expression levels of either protein via immunoblotting (Figure S1A), immunofluorescent staining of muscle sections showed differential distribution patterns for Alix in the soleus, which is composed primarily of slow fibers (Type I), and the gastrocnemius region near the soleus, which also contains Type I fibers (Figure S1B). Ozz protein was present throughout the myofibers, while Alix was found predominantly beneath the sarcolemma and faintly inside the myofibers (Figure S1C). Hematoxylin and eosin (H&E)-stained cross sections of the *Alix^−/−^*soleus muscle revealed gross abnormalities that closely mimicked those in the *Ozz^−/−^* muscle, i.e., numerous small-caliber fibers and fibers with central nuclei (Figure S1D). At the ultrastructural level, misaligned sarcomeric units with altered streaming of the Z bands, which is characteristic of *Ozz^−/−^* myofibers, were also present in *Alix^−/−^*myofibers (Figure 1A). Our most surprising observation was a feature shared by both mouse models display a prominent mitochondrial phenotype that consisted of supernumerary mitochondria of abnormal shape and size (Figure 1B).

**Figure 1.**
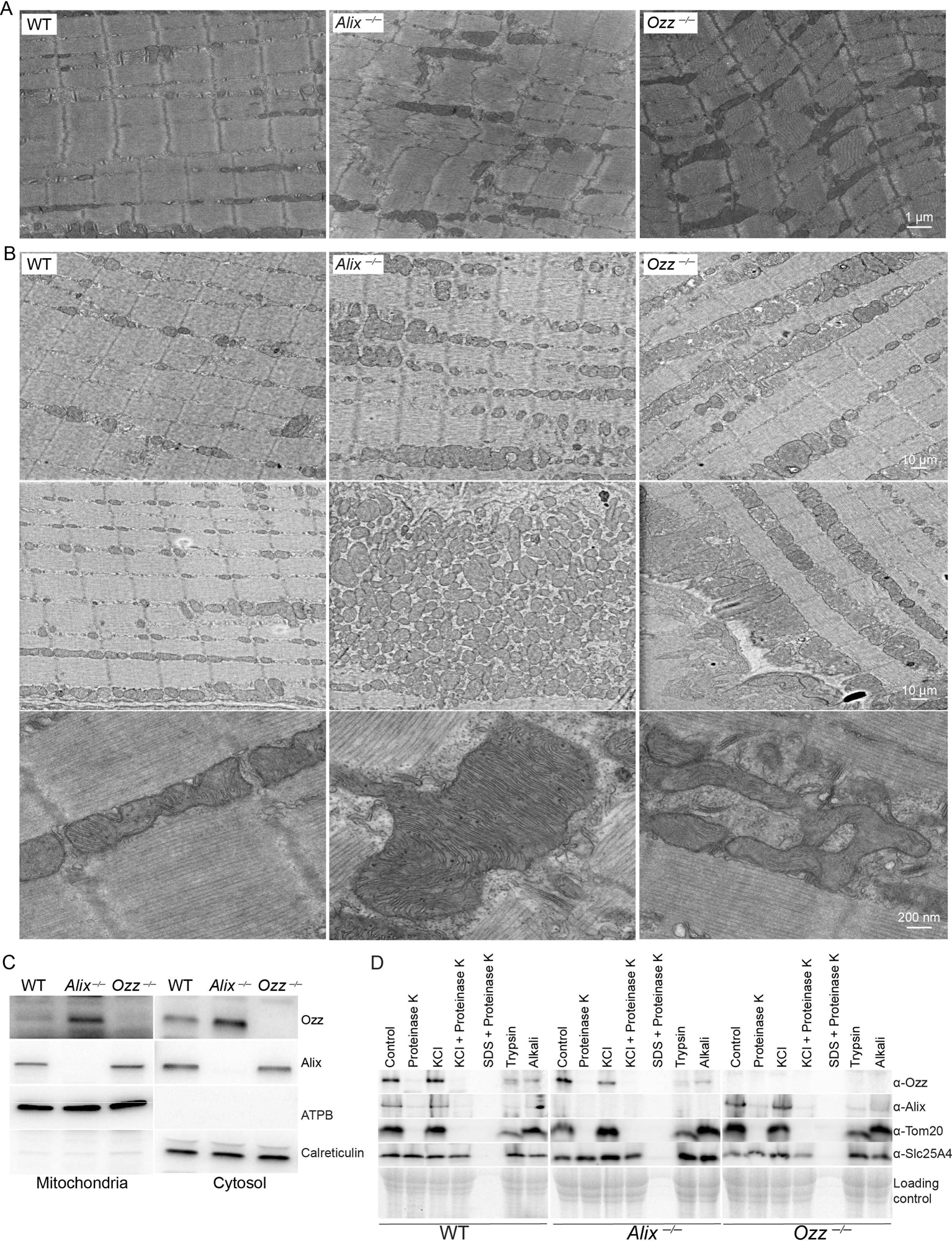
Abnormal myofibril organization and mitochondria in *Alix^−/−^* and *Ozz^−/−^* mice. (A) Transmission electron microscopy (TEM) of soleus muscle dissected from *Alix^−/−^* and *Ozz^−/−^*mice revealed a misalignment of sarcomeres with dysmorphic mitochondria compared to WT. Scale bar: 1 μm. (B) TEM of soleus muscle dissected from *Alix^−/−^*, *Ozz^−/−^*, and WT mice revealed abnormal mitochondrial morphology, size, and number. Scale bar: 10 μm or 200 nm. (C) Immunoblot analyses of the mitochondrial (2 μg) and cytosolic (6 μg) fractions of *Alix^−/−^*, *Ozz^−/−^*, and WT soleus muscles revealed endogenous Ozz and Alix. The purity of the subcellular fractionation of membranes & organelles was verified by immunoblotting using antibodies against a mitochondrial fraction marker (anti-ATPB) and an endoplasmic reticulum marker (anti-calreticulin). (D) Purified mitochondria were subjected to protease digestion before or after high-salt treatment, followed by immunoblot analyzes with anti-Ozz, anti-Alix, anti-Tom20 (OMM), and anti-Slc25A4(IMM) antibodies.

These findings led us to test if Ozz and Alix were localized to mitochondria. We detected both proteins in purified mitochondrial preparations from wild-type (WT) muscle tissue, but in those preparations from *Alix*^−/−^ muscles and *Ozz*^−/−^ muscles, the levels of each protein appeared to increase in the absence of the other (Figure 1C), suggesting that these proteins reciprocally regulate each other in the mitochondria. We next examined the submitochondrial distribution of Ozz and Alix. Purified mitochondria from WT, *Alix^−/−^*, and *Ozz*^−/−^ skeletal muscles were subjected to protease treatment and alkaline extraction, followed by immunoblotting with specific antibodies. Upon treatment with proteinase K, Ozz and Alix were degraded (Figure 1D), as was Tom20, a component of the OMM. Slc25A4, which resides in the IMM, was not affected (Figure 1D). All proteins, including Slc25A4, remained intact in mitochondria subjected to osmotic shock with potassium chloride (KCl). Combining KCl and proteinase K degraded Ozz, Alix, and Tom20 but not Slc25A4, whose levels were only slighted reduced (Figure 1D). Treatment with SDS (sodium dodecyl sulfate) and proteinase K degraded all proteins, irrespective of their localization. These results indicated that Ozz and Alix localize to the OMM. To further determine whether both proteins are associated with the luminal or cytosolic side of the OMM or are embedded in the OMM, we analyzed their sensitivity to trypsin digestion or alkaline pH treatment (Figure 1D). Incubation of purified mitochondria with trypsin resulted in the complete digestion of Ozz, Alix, and Tom 20 (Figure 1D). In contrast, alkaline treatment removed Ozz and Alix from the OMM, but it did not remove Tom20 (Figure 1D). Thus, Ozz and Alix are associated with the OMM, topologically face the cytosolic side, and are not inserted into the membrane. These findings provided the first indication that like Ozz, Alix may establish and/or maintain the sarcomeric apparatus and help preserve mitochondrial homeostasis.

### Ozz contains a mitochondrial targeting sequence

To determine how Ozz and/or Alix would traffic to the mitochondria, we examined the primary structures of both proteins and identified a putative mitochondrial-targeting sequence present only in Ozz (Mitoprot software). The sequence consisted of 36 amino acids that overlapped with the first NHR domain (Figure 2A). To validate this finding, we used CRISPR/Cas9 to engineer mutant mice that carried sequential nucleotide deletions, resulting in the loss of two, five, or nine amino acids (Δ2aa, Δ5aa, and Δ9aa) at the N terminus of the sequence (Figure 2A). Immunoblots of mitochondrial preparations from the muscles of these mutant mice showed that deletion of five or nine amino acids from the sequence was sufficient to completely abrogate Ozz localization to the mitochondria (Figure 2B); the Δ2aa mutant still allowed for trace amounts of Ozz to reach the organelles (Figure 2B). These results confirmed that this sequence motif in Ozz is responsible for its mitochondrial localization. One of the CRISPR/Cas9-engineered mutant mice, *ccOzz*^−/−^, carried a single base pair deletion that introduced a stop codon eight amino acids downstream from the start codon and truncated almost the entire open reading frame of *Ozz* (Figure 2A). Thus, *ccOzz*^−/−^ mice served as proper controls for all CRISPR/Cas9-derived mutants.

**Figure 2.**
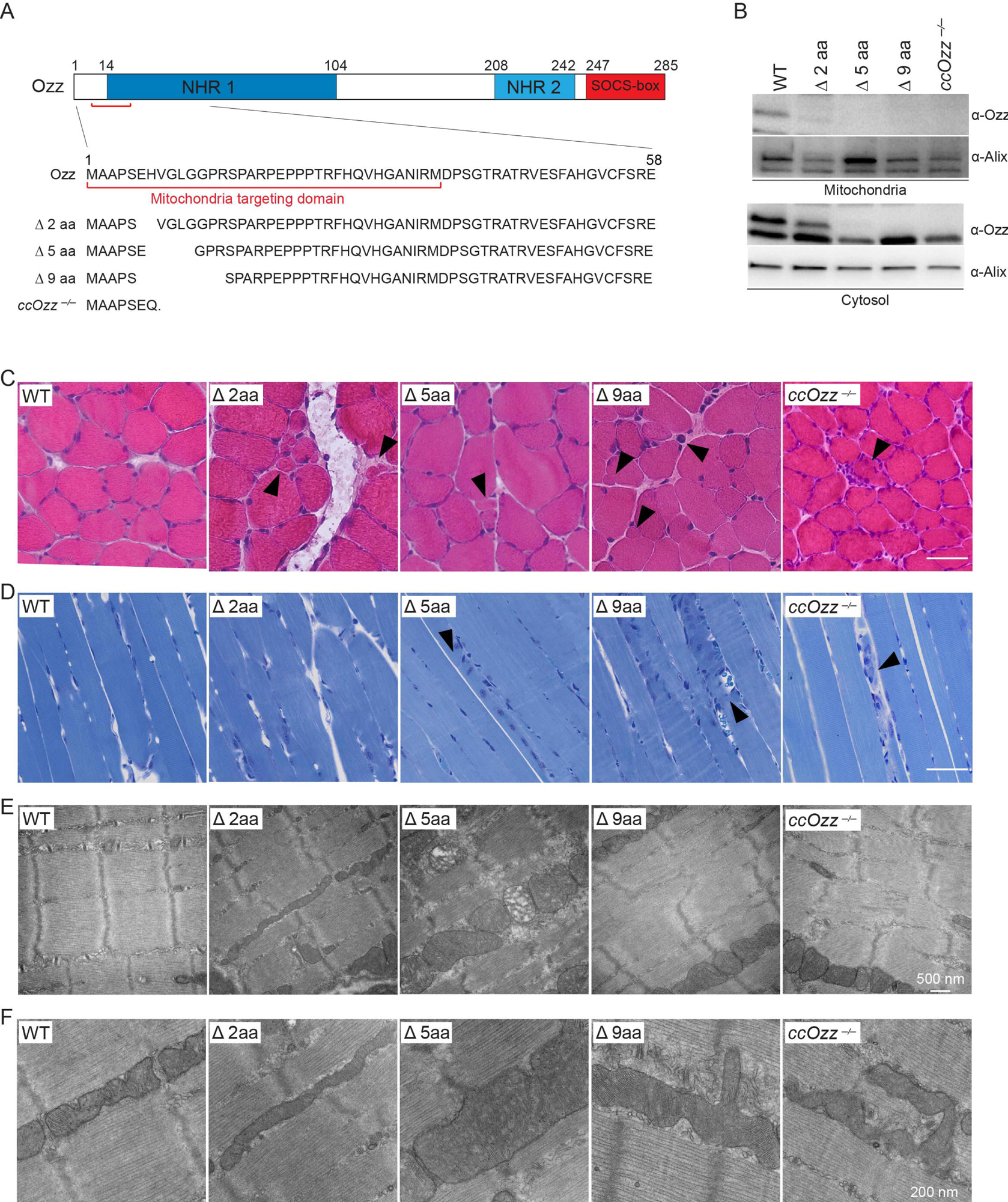
Ozz protein contains a mitochondrial-localization sequence. (A) Identification of a mitochondrial-targeting sequence in the Ozz protein’s primary structure. The numbers refer to amino acid (aa) positions, starting from the N-terminal end. Alignment of the Ozz aa sequence of mutant mice with that of WT mice confirmed the deletion of 2, 5, and 9 aa in exon 1 of the Ozz protein generated by the CRISPR/Cas9 system. (B) Immunoblot analyses of purified mitochondria probed with anti-Ozz antibody showed the absence of Ozz in Δ5aa, Δ9aa, and *ccOzz^−/−^* muscle, whereas the Δ2aa muscle showed trace amounts of Ozz. (C) H&E-stained cross sections of soleus muscle from 2-month-old WT, Δ2aa, Δ5aa, Δ9aa, and *ccOzz^−/−^* mice showed an increased frequency of small fibers (arrowheads) in CRISPR/Cas9 Ozz-deletion mutant mice compared to WT littermates. Scale bar: 50 μm (D) Toluidine blue staining of semi-thin sections of soleus muscles of 2-month-old WT, Δ2aa, Δ5aa, Δ9aa, and *ccOzz^−/−^* mice displayed fibers with central nuclei (black arrowheads). Scale bar: 50 μm. (E) TEM of the soleus muscle isolated from 2-month-old WT, Δ2aa, Δ5aa, Δ9aa, and *ccOzz^−/−^* mice. The fibers of mutant mice showed disorganization of the myofibrillar array associated with the abnormal appearance of the Z bands and surrounding structures. Scale bar: 500 nm. (F) TEM of the soleus muscle isolated from 2-month-old WT, Δ2aa, Δ5aa, Δ9aa, and *ccOzz^−/−^* mice revealed abnormal mitochondrial morphology, size, and number. Scale bar: 200 nm

Histopathologic examination of the soleus muscles isolated from all deletion mutants revealed gross morphologic defects that were consistent with those previously described in *Ozz^−/−^* mice ^31^. Numerous small-caliber fibers, an abnormal striation pattern, and an increased number of immature fibers with central nuclei were evident in H&E (Figure 2C) and toluidine blue-stained muscle sections from the Δ5aa, Δ9aa, and *ccOzz*^−/−^ mice (Figure 2D). The Δ2aa mice appeared to be the least affected (Figure 2C); however, at the ultrastructural level, all mutant mice, including the Δ2aa, showed the same myofiber abnormalities detected in the *Alix*^−/−^ and *Ozz*^−/−^ models (Figure 2E), i.e., misalignment of the sarcomeres, abnormal streaming of the Z-band, and defective mitochondria (Figure 2F).

### Mitochondrial respiration is impaired in *Alix*^−/−^and *Ozz*^−/−^ skeletal muscle

Deficiency of Ozz or Alix appeared to affect the mitochondrial morphology in a similar way. However, to better characterize this phenotype in both models, we needed to determine if the alterations affected mitochondrial function to the same extent. We, therefore, measured the oxygen consumption rate (OCR), which indicates the functional bioenergetic capacity of mitochondria, in cultures of primary proliferating myoblasts (Day 0) and differentiated myocytes/myotubes (Day 3) isolated from *Alix^−/−^* and *ccOzz*^−/−^ limb muscles (Figure 3A and 3B). Although the OCR profiles of myoblasts and myotubes were similar, the profiles of the *Alix*^−/−^and *ccOzz*^−/−^ Day 3 samples were significantly lower than those of WT controls: basal mitochondrial respiration normalized by subtracting nonmitochondrial OCR was decreased two folds in *Alix*^−/−^ myotubes and three folds in *ccOzz*^−/−^ myotubes compared to the WT control (*p* = 0.0001; Figure 3B). Oligomycin A inhibition showed similar proton leak results in all cells tested, indicating that most of the basal mitochondrial respiration in WT cells and mutant cells was devoted to ATP production (Figure 3B). Treatment of WT myotubes and mutant myotubes with the mitochondrial-uncoupling agent FCCP, which directly transports protons across the IMM instead of through the ATP synthase proton channel, enabled us to measure maximum oxygen consumption (Figure 3B). This approach revealed an even greater difference in OCR between *Alix*^−/−^*, ccOzz*^−/−^, and WT control myotubes (*p* <0.0001; Figure 3C). The *ccOzz*^−/−^ (*p* =0.0001) myotubes had the lowest respiratory reserve capacity, and *Alix*^−/−^ (*p* <0.0006) myotubes had values intermediate between *ccOzz^−/−^*cells and WT controls (Figure 3B), suggesting that Ozz is more important in maintaining mitochondrial respiratory capacity than is Alix. Comparable OCR profiles were obtained in proliferating *Alix*^−/−^ and *ccOzz*^−/−^ Day 0 myoblasts (Figure S2A and S2B).

**Figure 3.**
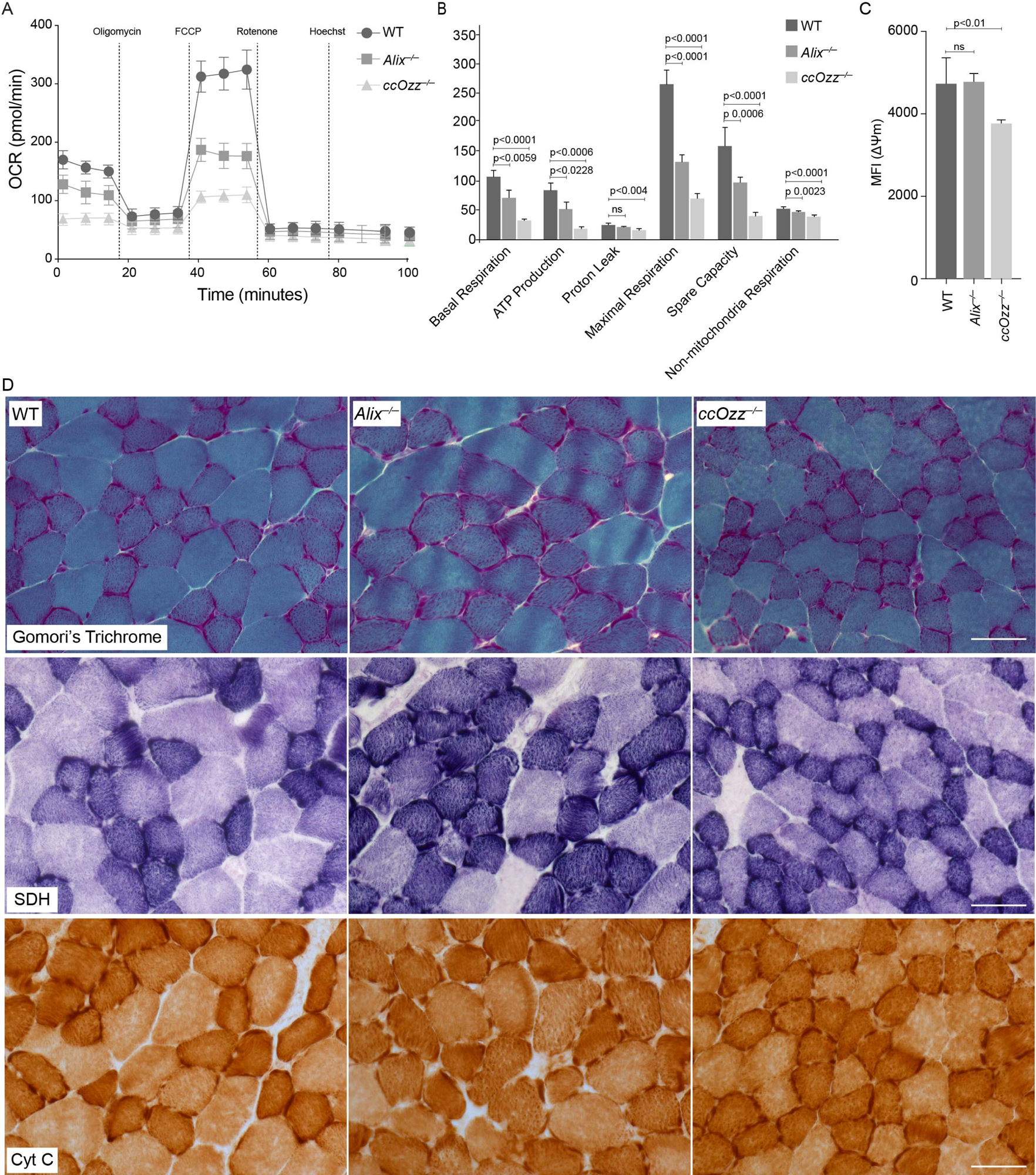
Analysis of mitochondrial phenotype in *Alix^−/−^* and *ccOzz^−/−^* skeletal muscle. (A) Oxygen consumption rate (OCR) profile of primary myotubes of WT, *Alix^−/−^*, and *ccOzz^−/−^*mice (Day 3) in response to treatment with oligomycin (ATP-linked respiration and proton leak), FCCP (mitochondrial reserve capacity), and rotenone/antimycin A (nonmitochondrial respiration). Hoechst 33342 staining was used to count the number of cells. (B) Analysis of the different stages of myoblast respiration determined from the OCR analysis of WT, *Alix^−/−^*, and *ccOzz^−/−^*primary myotubes (Day 3). Data are presented as mean ± SD, Student’s (unpaired) *t*-test; n = 3 independent samples. (C) Quantitative analysis of ΔΨM by using the mean fluorescence intensity (MFI) of TMRE in the primary myotubes of WT, *Alix^−/−^*, and *ccOzz^−/−^* muscle. Data are presented as mean ± SD, Student’s (unpaired) *t*-test; n = 100 cells per n = 3 independent samples. (D) Modified Gomori trichrome, SDH, and Cyt C staining of WT, *Alix^−/−^*, and *ccOzz^−/−^*soleus muscle sections. Scale bars:50 μm.

To examine the effect of Alix or Ozz deficiency on the mitochondrial membrane potential (ΔΨm), we used the TMRE probe to measure ΔΨm during the proliferation and differentiation of *Alix*^−/−^ and *ccOzz*^−/−^ myoblasts. TMRE is a cell-permeant fluorescent dye that accumulates in healthy mitochondria with an intact ΔΨm. Upon dissipation of the potential, TMRE ceases to accumulate in mitochondria, and the reduced fluorescence intensity serves as an indicator of the loss of mitochondrial membrane integrity. We found that the mean TMRE fluorescence intensity in *ccOzz*^−/−^ myotubes was decreased ∼20% compared to that in WT myotubes (*p* < 0.01) (Figure 3C); however, we found no significant differences in ΔΨm between the *Alix^−/−^* myotubes and the WT control (Figure 3C). After exposure to FCCP, which was used as a control of ΔΨm, the fluorescence intensity of the labeled mitochondria completely decayed, confirming the specific targeting of the probe to mitochondria. The same analysis performed in proliferating myoblasts did not show significant differences across WT, *ccOzz*^−/−^, or *Alix^−/−^*cells (Figure S2C). Thus, the absence of Ozz from skeletal muscle mitochondria had a greater effect on the respiratory chain than did the absence of Alix.

We then sought to determine if the functional defects observed in the mitochondria of *ccOzz*^−/−^ and, to a lesser extent, *Alix*^−/−^ myotubes were paralleled by specific histochemical changes in the corresponding muscle tissue that usually occur in human mitochondrial myopathy^10, 12^. For this, we applied three standard mitochondrial staining methods (i.e., modified Gomori trichrome, succinate dehydrogenase [SDH] and cytochrome C [Cyt C]) to cross-sections of soleus muscle from adult WT, *Alix*^−/−^and *ccOzz*^−/−^ mice. In *Alix*^−/−^and *ccOzz*^−/−^ muscles, Gomori trichrome staining identified an increased number of deep, red-colored myofibers with ragged contours and a subsarcolemmal accumulation of the dye; these features indicated mitochondrial myopathy (Figure 3D). This phenotype was coupled with an increased number of fibers intensely stained with SDH and Cyt C, suggesting an overall enhanced enzymatic activity of mitochondria in the *ccOzz*^−/−^and *Alix*^−/−^ soleus muscles (Figure 3D).

### Mitochondrial dysfunction in *Alix^−/−^* and *ccOzz^−/−^*skeletal muscle results in mitophagy

Overall, the mitochondrial phenotypic alterations in *Alix^−/−^*and *ccOzz^−/−^* skeletal muscles further supported the notion that mitochondria are dysfunctional in both mutant myofibers. Given that defective mitochondria are removed by selective mitophagy under physiological conditions^45–47^, we tested whether this QC process was activated in the mutant mice. To perform such an analysis, we crossed *Alix*^−/−^ mice and *ccOzz*^−/−^ mice with the *mito*-QC reporter mouse model, which expresses a GFP-mCherry construct specifically targeted to mitochondria^48^. In these models, activation, and quantification of mitophagy could readily be assessed by measuring the number of red-only puncta, representing autophagosomes with engulfed mitochondria that fused to acidic lysosomes, thereby quenching the GFP fluorescence signal^48^. Confocal microscopy analyses of the soleus muscles from these crossings revealed a significant increase of red-only puncta in both *mito-*QC/*Alix*^−/−^ (100) and *mito-*QC*/ccOzz*^−/−^ (98) mice compared to WT (30) controls (Figures 4A and S3). These data showed that the mitochondrial dysfunction associated with ablating Ozz or Alix induced mitophagy in skeletal muscle.

**Figure 4.**
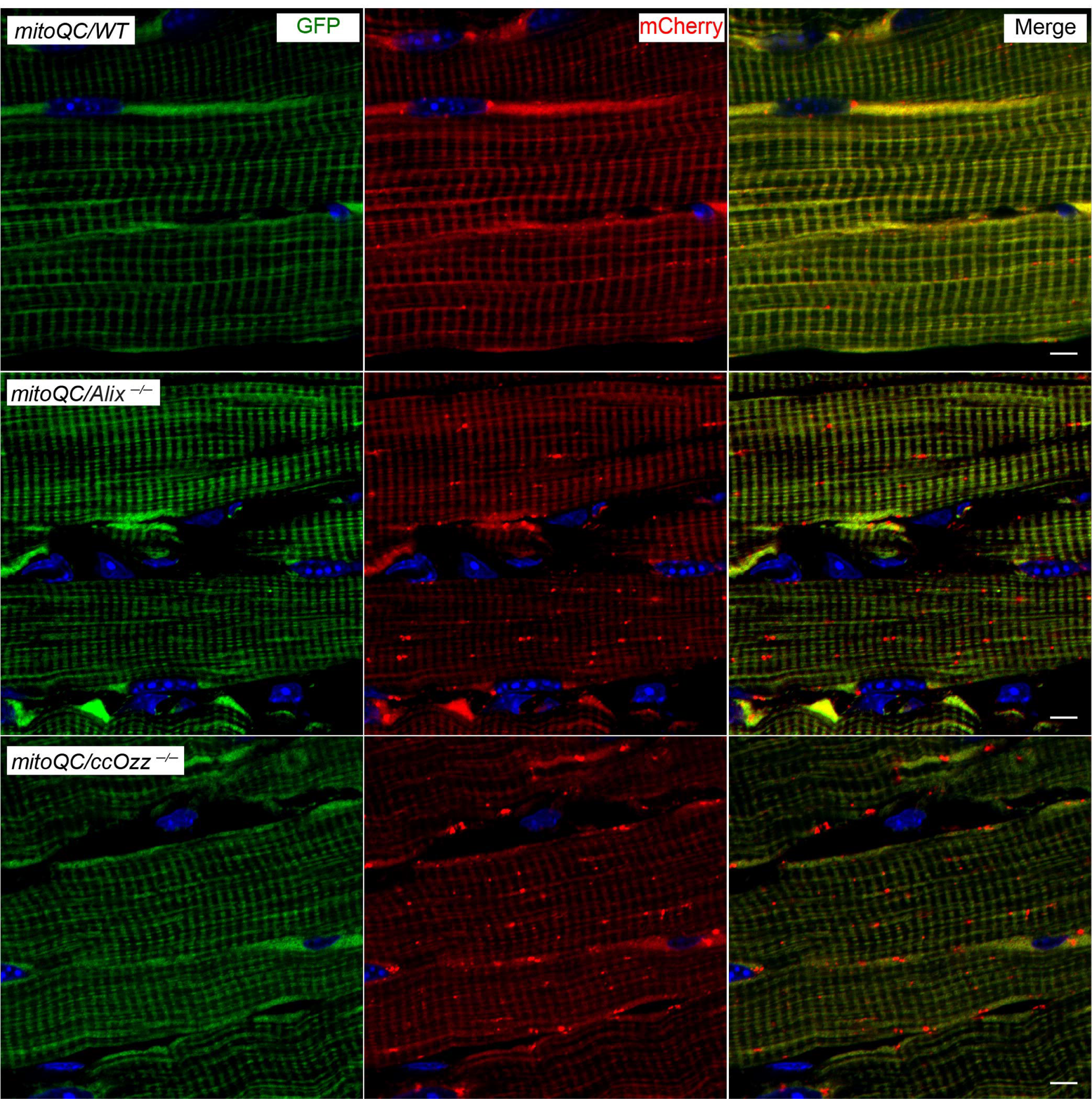
Mitophagy analysis using MitoQC reporter mice. Representative confocal images of the soleus muscle isolated from MitoQC/WT, MitoQC/*Alix^−/−^* and MitoQC/cc*Ozz^−/−^* mice show increased mitophagy (red) in both MitoQC/*Alix^−/−^* and MitoQC/cc*Ozz^−/−^* mice. Scale bars: 5 μm.

### Metabolomics of *Alix*^−/−^ and *ccOzz*^−/−^ soleus muscles underscore their mitochondrial abnormalities

Our data revealed that *Alix*^−/−^ and *ccOzz*^−/−^ myofibers had both functional and biochemical abnormalities in their mitochondria that could reflect an overall impaired mitochondrial metabolism. We investigated this possibility by performing high-throughput metabolomic analyses of *Alix*^−/−^, *cc*Ozz^−/−^, and WT soleus muscles isolated from 8-week-old mice. We detected ∼120 metabolites in the *Alix*^−/−^ soleus (Table S1) and ∼100 metabolites in the *ccOzz*^−/−^ soleus (Table S2) that showed significantly different levels than the corresponding metabolites in WT controls; 41 unique metabolites were altered in both *Alix^−/−^* and *ccOzz^−/−^* soleus muscles (Figure 5A, Table S3). Using MetaboAnalyst 4.0 software, we generated heat maps of differentially expressed metabolites in each mutant muscle compared to WT controls (Figure 5A and 5B). Combining the heat map values for each metabolite revealed distinct metabolic profiles for *Alix^−/−^*, *ccOzz^−/−^*, and WT soleus muscles (Figure 5B). Even more striking was the finding that several metabolites (i.e., ADP, CoA, Oxa, thymidine, pyruvate, UDP, and UTP) had opposite expression levels in *Ozz*^−/−^ and *Alix*^−/−^ samples that still differed from the WT control. Interestingly, many of them (i.e., ADP, CoA, citrate, dUMP, and malate) appeared to be regulated by the mitochondrial SLC25 solute carrier family of proteins^49–52^. Pathway analyses of the differentially expressed metabolites in *Alix*^−/−^, *ccOzz*^−/−^, and WT muscles enabled us to link the metabolites to specific metabolic pathways, such as glycolysis/glucogenesis, starch/sucrose metabolism, pyrimidine metabolism, citric acid cycle, lactose synthesis, and transfer of acetyl groups to mitochondria (Figure 5C and 5D, Tables S4 and S5). Metabolites belonging to the same pathway were equally affected in *Alix*^−/−^ and *ccOzz*^−/−^ muscles. Specifically, glycolysis/glucogenesis was the dominant pathway upregulated in the *ccOzz*^−/−^ muscle, compared to that in the *Alix*^−/−^ and WT muscles (Figure 5C and 5D, Tables S4 and S5).

**Figure 5.**
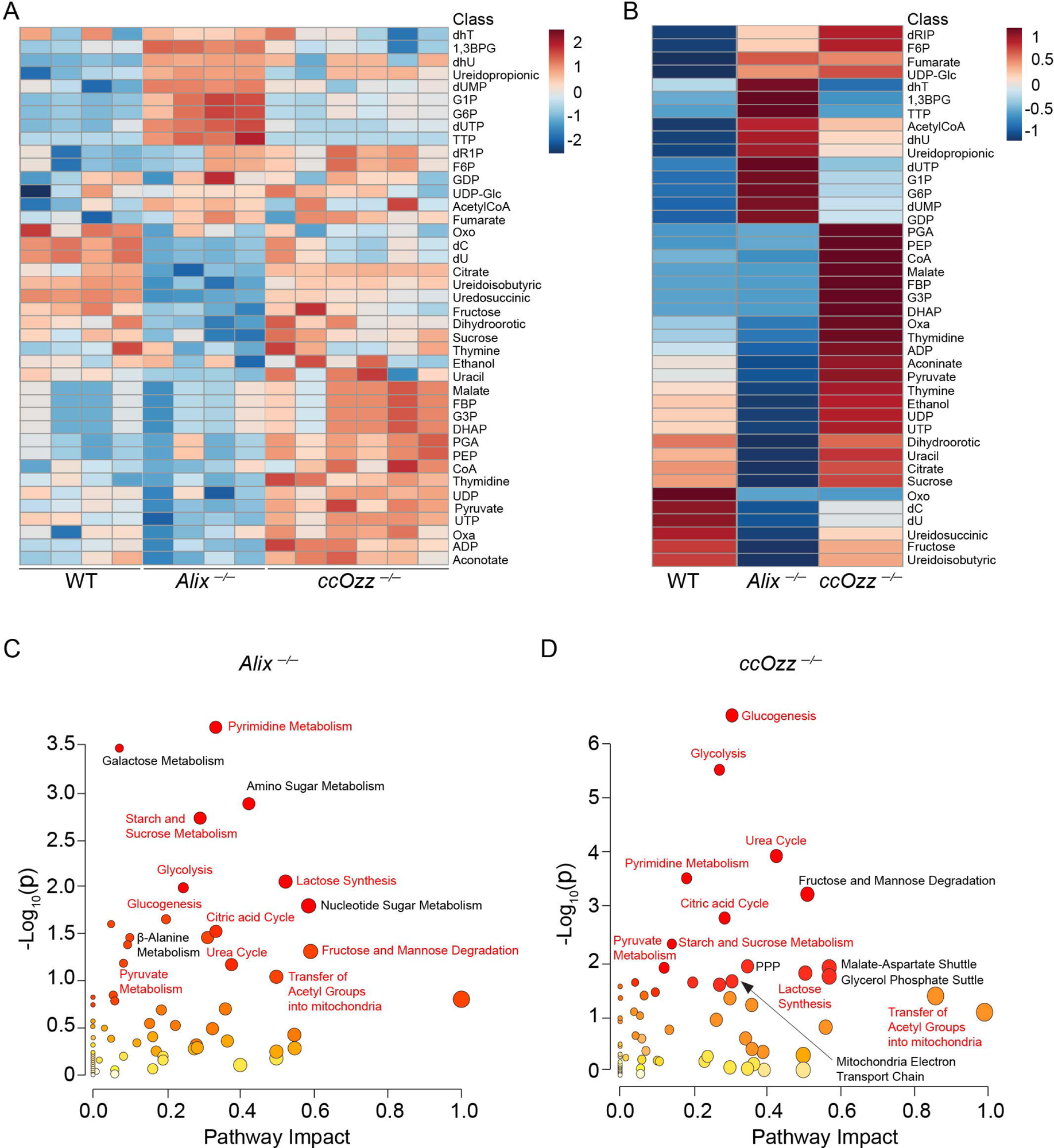
Enrichment of metabolic pathways in *Alix^−/−^* and *ccOzz^−/−^* soleus skeletal muscle. (A) Clustered heat map of metabolites detected at significantly different (p <0.05) levels between *Alix^−/−^*(n = 4) or *ccOzz^−/−^* (n = 6) soleus muscles and WT control (n = 4) soleus muscle samples. (B) Clustered heat map (grouped) of the top 41 metabolites detected at significantly different (p <0.05) levels between *Alix^−/−^* (n = 4) or *Ozz^−/−^* (n = 6) soleus muscles and WT control (n = 4) soleus muscle samples. (C) Functional enrichment and pathway topology analysis of metabolites affected in *Alix^−/−^* soleus muscle compared to WT control by using the MetaboAnalyst 5.0. Pathway Analysis tool. (D) Functional enrichment and pathway topology analysis of metabolites affected in *ccOzz^−/−^* soleus muscle compared to WT control by using the MetaboAnalyst 5.0. Joint Pathway Analysis tool.

### Ablation of Ozz or Alix in vivo induces a muscle fiber switch

To further investigate the altered mitochondrial metabolism in the skeletal muscles of both mutant mice, we examined the relation between fiber-type composition and fiber-energy metabolism in the soleus muscle. The soleus muscle is specified by a high percentage of fibers expressing myosin heavy chain-beta (MyHC-β) protein, which represents slow oxidative Type I fibers, and MyHC-IIA proteins, which represent fast oxidative/glycolytic Type IIA fibers. We analyzed MyHC expression levels in serial cross sections of soleus muscles isolated from *Alix^−/^*^−^, *ccOzz*^−/−^, and WT mice by immunofluorescent staining of the MyHC isoforms Type I, Type IIA, and Type IIB. By applying an ImageJ macro program, we separated the fiber types by their fluorescent staining pattern and accurately quantified each type. The WT soleus muscle contained the expected percentages of slow Type I fibers (46%) and fast Type IIA fibers (54%) (Figure 6A); however, the number of Type IIB fibers was too low to be considered in this analysis. The *Alix*^−/−^ soleus muscle contained 42% slow Type I fibers and 58% fast Type IIA fibers, and the Ozz^−/−^ soleus muscle contained only 32% slow Type I fibers and 68% fast Type IIA fibers (Figure 6B). Thus, both mutants had an increased percentage of glycolytic/oxidative Type IIA fibers at the expense of oxidative Type I fibers in their soleus muscles, but the deficiency of Ozz was more pronounced, with about a 30% decrease in Type I fibers versus only ∼9% in the *Alix*^−/−^ muscle. Thus, ablation of Ozz or Alix affected the myofiber composition of the soleus muscle.

**Figure 6.**
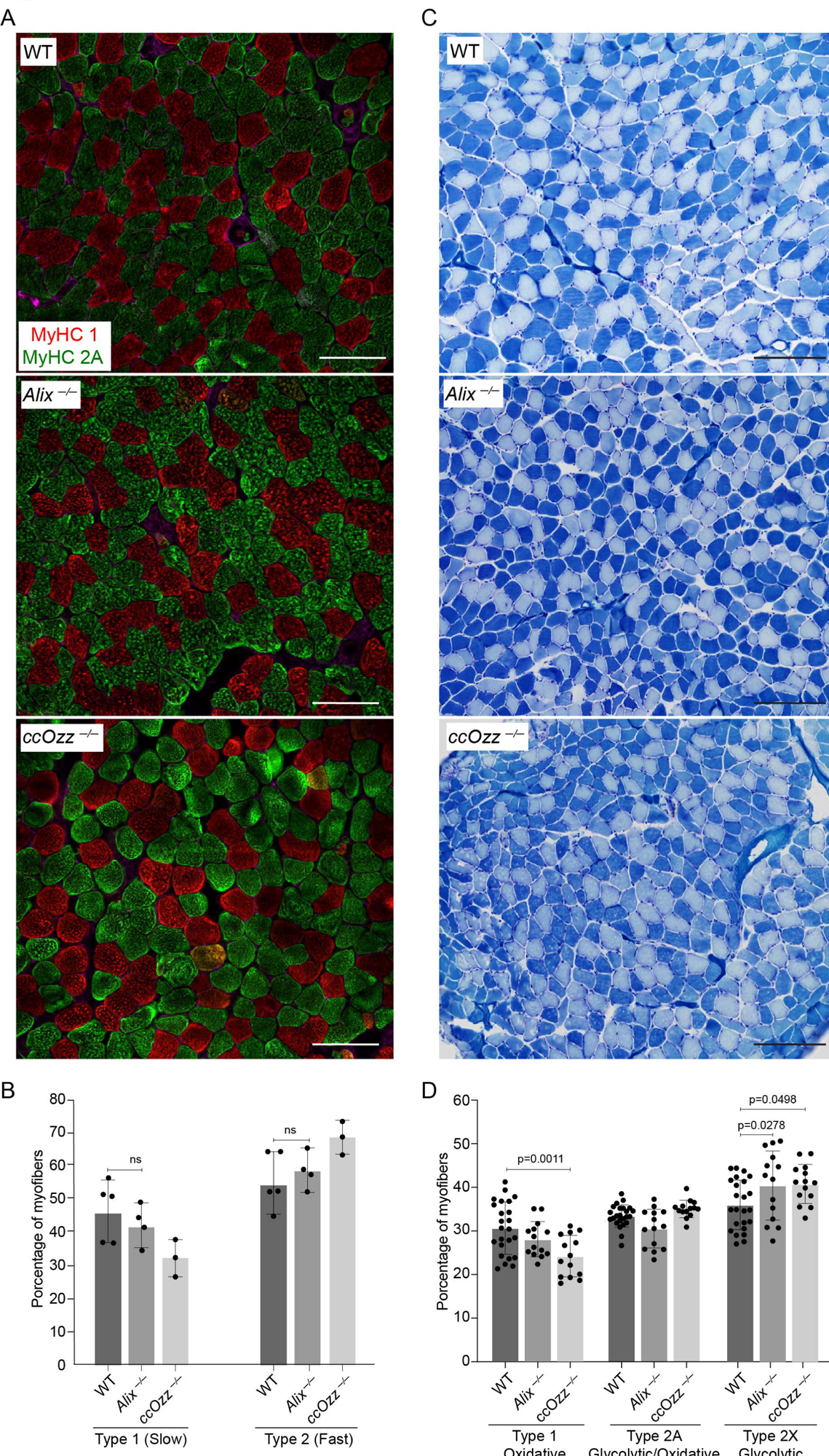
Fiber-type switching in muscles of *Alix^−/−^* and *ccOzz^−/−^* mice. (A) Immunofluorescence staining of soleus muscle sections from WT, *Alix^−/−^*, and *ccOzz^−/−^*mice with antibodies specific for MyHC Type I (red) or MyHC Type IIA (green) fibers. Scale bar: 100 μm. (B) Quantification of the average percentage of Type I and Type IIA fibers from panel A. Data are presented as mean ± SD, Student’s (unpaired) *t*-test; n = 4 independent samples. (C) Representative images of metachromatic ATPase staining on cross sections of soleus muscle from WT, *Alix^−/−^*, and *ccOzz^−/−^* mice. ATPase activity stains Type I fibers (dark blue), Type IIA fibers (light blue), and Type IIB fibers (very light blue). Scale bar: 200 μm (D) Quantification of the average percentage of Type I (oxidative), Type IIA (glycolytic/oxidative) and Type IIX (glycolytic) fibers from panel C. Data presented are the mean ± SD, Student’s (unpaired) *t*-test; n = 4 independent samples.

We then asked whether the myosin immunofluorescence staining pattern reflected the functional ATPase activity of individual myofibers. For this experiment, we applied a metachromatic staining of myofibrillar ATPase approach to 12-µm-thick frozen serial sections of soleus muscles from *Alix^−/−^*, *ccOzz*^−/−^, and WT mice. Following incubation of the sections with ATP in the presence of calcium, we distinguished three types of fibers based on their color intensity: Type I oxidative, light blue; Type IIA glycolytic/oxidative, intermediate blue; and Type IIX glycolytic, dark blue (Figure 6C). Using a similar ImageJ macro program, we set thresholds for the color intensity to separate and quantify the fiber types. The WT soleus muscle consisted of a mixture of approximately 31% oxidative Type I fibers, 33% glycolytic/oxidative Type IIA fibers, and 36% glycolytic Type IIX fibers (Figure 6D). In contrast, the *Alix^−/−^*soleus contained about 28% oxidative Type I fibers, 31% glycolytic/oxidative Type IIA, but had a comparable percentage (∼36%) of glycolytic Type IIX fibers. Once again, the *ccOzz^−/−^* soleus muscle was the most affected, containing only ∼24% oxidative Type I fibers, and increased percentages of glycolytic/oxidative Type IIA (35%) and glycolytic Type IIX (41%) fibers.

These functional analyses showed that the deficiency of Alix or Ozz changed the myofiber composition of the soleus muscle, which was consistent with our OCR results, and this change affected the overall mitochondrial metabolism of this muscle type.

### CRL5^Ozz^ in mitochondria targets a SCL25 solute carrier family member for proteasomal degradation

Having established the mitochondrial localization of Ozz, we wanted to ascertain whether the other components of CRL5^Ozz^ also colocalized in this organelle to define a role for this ligase and its substrate Alix in mitochondria. We first tested whether EloC, EloB, Rbx1, and Cul5 could be detected by coimmunoprecipitation and/or immunoblotting of mitochondria isolated from WT, *Alix^−/−^*, or *ccOzz*^−/−^ skeletal muscle. These initial experiments were unsuccessful, probably because Ozz and Alix are expressed at rather low levels in steady-state mitochondria. To overcome this caveat, we overexpressed recombinant Ozz fused to a short peptide tag (Avi-tag) that was biotinylated by the coexpressed bacterial ligase BirA^53, 54^. Cells transduced with an empty vector were used as controls. We incubated lysates of mock-transduced and Avi-Ozz– overexpressing, differentiated C2C12 myotubes (Day 3) at the peak of Ozz expression ^31^ with streptavidin beads to capture biotinylated Avi-Ozz and any of its interactors. Immunoblot analysis of the pulled-down proteins from lysates of overexpressing myotubes demonstrated that Avi-Ozz was associated with all other components of the CRL5^Ozz^ ligase and with Alix, indicating that biotinylated Avi-Ozz assembled into an active E3 ligase (Figure 7A). Aliquots of the streptavidin-captured proteins were also subjected to mass spectrometric analyses, which confirmed the pull-down of Ozz’s direct interactors EloB and EloC (Table S6). However, the most surprising finding was that several members of the mitochondrial solute carrier family, including the muscle specific Slc25A4, were pulled down with Ozz and could be potential substrates of CRL5^Ozz^ (Table S6).

**Figure 7.**
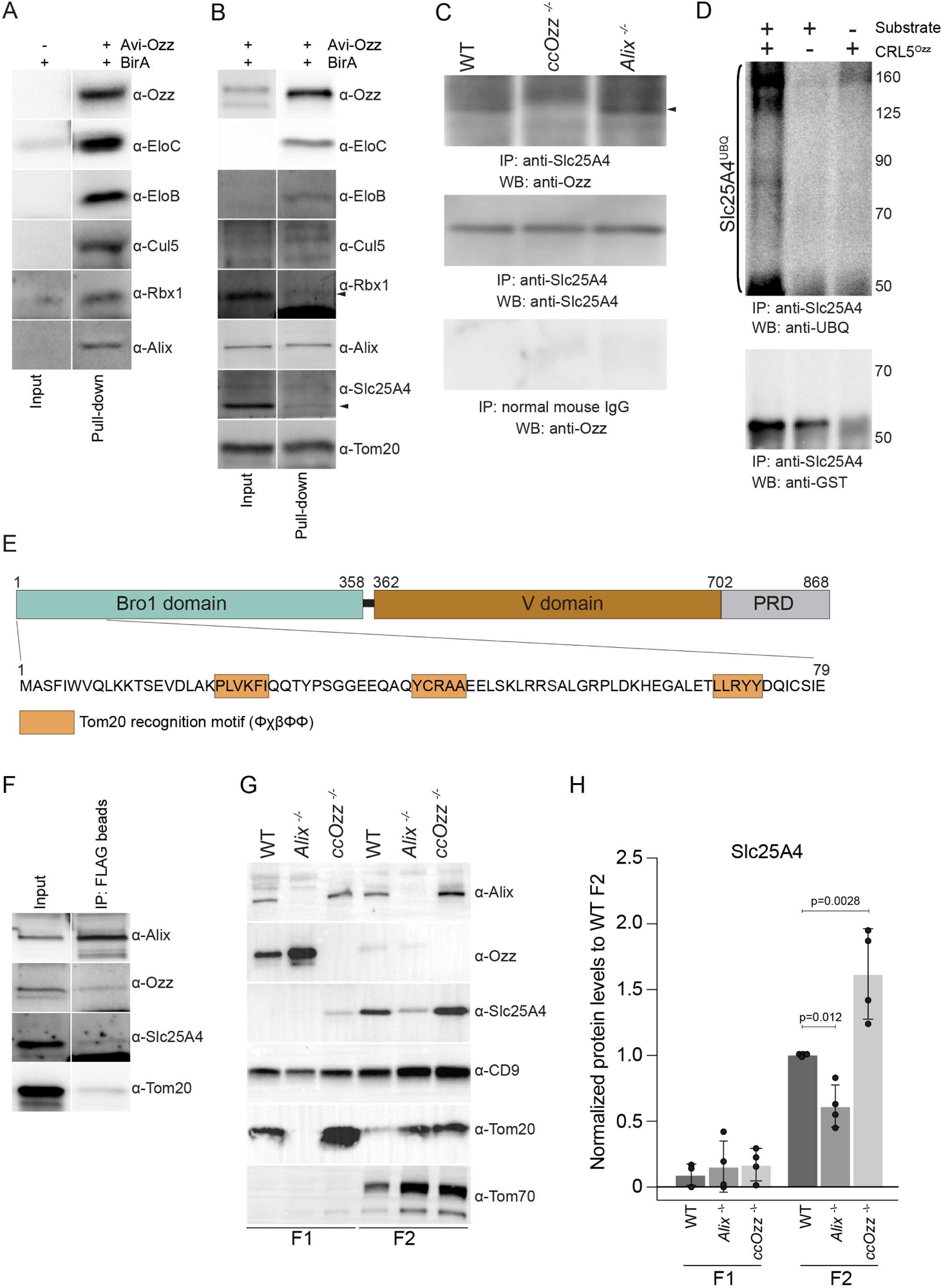
Alix and Ozz cooperatively regulate Slc25A4. (A) Immunoblot analyses of biotinylated cytosolic Avi-Ozz from C2C12 myotubes (Day 3) probed with anti-Ozz, anti-EloC, anti-EloB, anti-Cul5, anti-Rbx1, and anti-Alix antibodies. (B) Biotinylated mitochondrial Avi-Ozz from preparations of C2C12 myotubes (Day 3) were analyzed by Western blot using antibodies to detect CRL5^Ozz^ ligase components (EloB, EloC, Cul5, Rbx1), Alix substrates/interactors, Slc25A4, Tom20, and Avi-Ozz protein. (C) Immunoprecipitation (IP) of mitochondrial lysates isolated from WT, *Alix^−/−^*, and *ccOzz^−/−^*muscles showed co-precipitation of Slc25A4 and Ozz. Immunoblots were probed with anti-Ozz and anti-Slc25A4 antibodies. IgG was used as a loading control. (D) In vitro ubiquitination of GST-Slc25A4, followed by IP with anti-Slc25A4 antibody and probed with anti-ubiquitin (UBQ) and anti-Slc25A4 antibodies showed the interaction of Slc25A4 with the CRL5^Ozz^ complex. (E) Protein domain mapping of Tom20-recognition motifs (orange boxes) identified in the Alix primary protein structure. The numbers refer to the amino acid positions starting from the N-terminal sequence of Alix protein. (F) IP of purified FLAG-tagged Alix from C2C12 myotubes (Day 3) probed with anti-Alix, anti-Ozz, anti-Slc25A4, and anti-Tom20 antibodies showed the interaction of Slc25A4 with Alix. (G) Immunoblot analyses of pooled sucrose-gradient fractions of exosomes (F1: fractions 1-2 and F2: fractions 3-6) isolated from WT, *Alix^−/−^*, and *ccOzz^−/−^* muscles probed with antibodies against Alix, Ozz, Slc25A4, CD9, Tom20, and Tomm70. (H) Quantification of Slc25A4 from the pooled F1 and F2 fractions of soleus muscle. Data presented are the mean ± SD, Student’s (unpaired) *t*-test; n = 4 independent samples.

We then tested whether we could resolve an assembled CRL5^Ozz^ in mitochondria isolated from differentiated C2C12 myocytes overexpressing biotinylated Avi-Ozz (Figure 7B). For this, we subjected mock-transduced cells and Avi-Ozz–overexpressing cells to differential centrifugation to obtain crude mitochondrial fractions. Immunoblots of these preparations showed the presence of the mitochondrial markers Tom20 and Slc25A4 (Figure 7B). Furthermore, Ozz, Alix, and Rbx1 were detected in Avi-Ozz–overexpressing cells; EloB, EloC, and Cul5 were barely visible. However, after streptavidin pull-down of biotinylated Avi-Ozz from these crude mitochondria, we detected all components of CRL5^Ozz^ (EloB, EloC, Cul5, and Rbx1), Alix, and Slc25A4 (Figure 7B), confirming the assembly of this ligase in mitochondria. The fact that Tom20, a member of the TOM complex, was also pulled down with Ozz (Figure 7B), led us to speculate that the localization of Ozz to the mitochondria might be regulated by direct or indirect interaction with the TOM complex. To verify that Slc25A4 interacts with Ozz in vivo, we immunoprecipitated lysates of mitochondria purified from gastrocnemius and soleus muscles with anti-Slc25A4 antibodies and on immunoblots probed them with anti-Ozz antibody (Figure 7C). Ozz co-immunoprecipitated with Slc25A4 from WT and *Alix*^−/−^ mitochondrial lysates (Figure 7C), indicating that these endogenous proteins were physically associated (Figure 7B and 7C).

To ascertain that Slc25A4 is a substrate of CRL5^Ozz^, we also performed in vitro ubiquitination assays using a purified, reconstituted CRL5^Ozz^ complex^30, 31^ and a GST-tagged full-length recombinant Slc25A4 (Figure 7D). The ubiquitinated products were then immunoprecipitated with anti-Slc25A4 antibody and probed on immunoblots with anti-ubiquitin, and anti-GST antibodies (Figure 7D). This in vitro reaction resulted in the polyubiquitination of Slc25A4, confirming that it is a substrate of CRL5^Ozz^ in mitochondria (Figure 7D).

### Alix regulates the loading of mitochondrial Slc25A4 into exosomes

Given that Alix and Slc25A4 were pulled down with biotinylated Avi-Ozz from differentiated myotubes, we determined whether Alix also interacts with Slc25A4. We overexpressed FLAG-tagged full-length Alix in C2C12 cells, followed by co-immunoprecipitation of Alix from the mitochondrial fractions of myotubes with anti-FLAG antibody and immunoblotting with antibodies against Ozz, Slc25A4, and Tom20 (Figure 7F). We also detected all three endogenous proteins in the crude mitochondrial fractions of cells overexpressing Alix (Figure 7F). Tom20 coimmunoprecipitated with flagged Alix, implying that these proteins interact directly via three Tom20-recognition motifs within its Bro1 domain (Figure 7E and 7F); however, this observation requires further investigation (Figure 7F). Furthermore, small amounts of Slc25A4 also coimmunoprecipitated with flagged Alix, suggesting that these two proteins interact in mitochondria.

Together, these findings hinted at a role of Alix in mitochondria. Based on the well-established function of this protein in cargo sorting and biogenesis of multivesicular body/exosomes, we analyzed the contents of extracellular vesicles/exosomes isolated from *Alix^−/^*^−^, *ccOzz*^−/−^, and WT gastrocnemius/soleus muscles. Purified vesicles were fractionated on sucrose density gradients, and the resultant fractions were probed on immunoblots with antibodies for a selected exosomal marker CD9, CD81 and mitochondrial markers Tom20 and Tom70 (Figures 7G and S4). The CD81-fractionation pattern showed that exosomes from all experimental samples sedimented in fractions 3-7 of the gradients at densities: 1.10 to 1.14 g/mL^55^ (Figure S4). Ozz, Alix, and Slc25A4 were detected in WT exosomes, though at low levels (Figure S4). As indicated by the sedimentation pattern of the F-actin control, protein levels in individual fractions were inconsistent due to difficult-to-control variability of the trichloroacetic acid (TCA) buffer precipitations. Thus, to resolve enough of the proteins of interest, we pooled fractions 1-2 (Pool 1) corresponding to heavy vesicles rich in endolysosomal markers, and fractions 3-7 (Pool 2) corresponding to the exosomal fractions (Figure 7G). We were unable to determine whether the exosomal preparations included MDVs, because it was experimentally difficult to isolate them from muscle mitochondrial preparations. However, immunoblot analysis of Pools 1 and 2 unequivocally demonstrated that not only Ozz and Alix but also Slc25A4 were recovered from exosomes in WT muscles (Figure 7G). As we previously observed in mitochondrial preparations, the absence of Ozz or Alix resulted in the accumulation of the other protein in both pools (Figure 7G). Intriguingly, Slc25A4 was decreased in *Alix^−/−^* exosomes but increased in *ccOzz^−/−^* exosomes (Figure 7G and 7H). Therefore, *ccOzz^−/−^*mitochondria may impair CRL5^Ozz^-mediated degradation of Slc25A4 and redirect the protein to exosomes. In contrast, Slc25A4 was not efficiently loaded into exosomes in *Alix^−/−^* mitochondria (Figure 7G and 7H).

## DISCUSSION

In this study, we uncovered a novel, essential role of the multimeric E3 ligase CRL5^Ozz^ in the mitochondrial function of skeletal muscle. We also discovered a previously unknown domain in Ozz, the substrate-binding component of CRL5^Ozz^, that is responsible for its mitochondrial localization. We showed that the loss of Ozz in skeletal muscle causes severe morphologic defects of mitochondria, which appear numerous, swollen, and dysmorphic. At the biochemical level, *ccOzz^−/−^* mitochondria have a decreased OCR and defective ΔΨm. Similar, albeit less severe, phenotypic alterations were observed in the absence of Alix. This combination of histopathologic, biochemical, and metabolic defects in the skeletal muscle of both mouse models, accompanied by increased mitophagy strongly suggests the occurrence of a mitochondrial myopathy and identifies Ozz and Alix as potential predisposing factors for these clinical conditions in humans.

We demonstrated that ablation of Ozz or Alix affects skeletal muscle metabolism and fiber composition. Mammalian skeletal muscle is composed of a mixture of muscle fibers, Type I and Type II (Type IIa, Type IIb, and Type IIx)^2^. Type I fibers, which are designed to maintain aerobic activity, contain numerous mitochondria and use predominantly OXPHOS for energy production; in contrast, Type II (IIx and IIb) fibers contain fewer mitochondria and more frequently depend on anaerobic glycolysis for energy production^2, 56^. Type IIa fibers have hybrid characteristics between Type I and Type IIb fibers, in that they have intermediate numbers of mitochondria and oxidative potential^2^. Muscle fibers are extremely plastic, and the fiber composition of muscles can change in response to various physiologic and/or pathologic conditions. Our data demonstrate that ablating Alix and Ozz in the soleus muscle promotes changes in MyHC isoform composition, leading to a switch from oxidative to glycolytic metabolism. To support this conclusion, we found that the metabolic profiles of the *Alix^−/−^* and *ccOzz^−/−^* soleus muscles differed significantly from that of their WT counterpart. This is supported by the elevated levels of metabolites associated with glycolytic myofibers in both mutant muscles, at the expense of those associated with oxidative myofibers. Most remarkably, we found an array of metabolites (e.g., ADP, pyruvate, uracyl, palmitate, oxoglutarate, and adenine) that are regulated by members of the SLC25 mitochondrial carrier family of proteins. These solute carriers mediate the transport of amino acids, carboxylic acids, fatty acids, cofactors, inorganic ions, and nucleotides across the IMM and are crucial for many cellular processes^49–52^, especially those regulating the oxidative capacity of the mitochondria.

We show for the first time that CRL5^Ozz^ localizes to the OMM and exposes part of its structure to the cytosol. At this location, the ligase controls the proteasomal degradation of Slc25A4, which we identified as a novel substrate of CRL5^Ozz^. We demonstrated that Alix^32^ also localizes to the mitochondria, where it regulates the loading of Slc25A4 into muscle-derived exosomes.

Mammalian Slc25A4, one of the most abundant isoforms among solute carrier proteins, is highly expressed in cardiac muscle and skeletal muscle^5^. In humans, SLC25A4 is present in small amounts in proliferating myoblasts, but its expression levels increase during myoblast differentiation^57^, a pattern of expression shared by Ozz. The *Slc25A4^−/−^* mouse model^58^ presents with abnormal, enlarged mitochondria, a characteristic of ragged-red fibers like those found in human mitochondrial myopathies and in *Alix^−/−^*and *ccOzz^−/−^* skeletal muscles. Biochemically, the mitochondria of *Slc25A4^−/−^* skeletal muscle exhibit defects in ADP-stimulated respiration that are consistent with impaired ATP/ADP transport across the IMM^58^. Contrary to Slc25A4 deficiency in mice, in vitro overexpression of this carrier saturates the mitochondrial import machinery in the cytosol, causing accumulation and aggregation of the protein. This triggers the so-called mitochondrial precursor overaccumulation stress, which is characterized by toxic accumulation of unimported mitochondrial proteins in the cytosol, increased protein ubiquitination, and apoptotic cell death^59, 60^. In humans, several missense mutations in the *SLC25A4* gene result in progressive external ophthalmoplegia, a multisystemic disease that manifests through weakness of the skeletal muscle^5, 61^. Therefore, tight regulation of SLC25A4 levels in mitochondria is required, as it is a predisposition factor for mitochondrial myopathies.

SLC25A4 is in the IMM and is required for transport of ADP into the mitochondrial matrix for the synthesis of ATP by OXPHOS. Newly synthesized mitochondrial ATP is then exported by the same carrier across the IMM in exchange for ADP. The selection of ATP versus ADP is controlled by the ΔΨm and the electrogenic capacity of SLC25A4. Neurons, cardiomyocytes, and hepatocytes, which rely on active aerobic mitochondrial metabolism, contain higher ATP/ADP ratios in the cytosol than in the mitochondrial matrix^62^. A high cytosolic ratio, downstream of aerobic mitochondrial metabolism, suppresses glycolysis^63–65^. However, when mitochondrial function is compromised, like in hypoxia/ischemia, the production of mitochondrial ATP decreases, the ATP/ADP ratio drops dramatically, and glycolysis is stimulated to generate ATP anaerobically. Thus, the cytosolic ATP/ADP ratio is a key feature that determines whether cell metabolism is predominantly oxidative or glycolytic. We now show for the first time that in skeletal muscle, Alix or Ozz affects mitochondrial ΔΨm, most likely due to a defective ATP/ADP ratio associated with abnormal turnover of the Slc25A4 carrier.

Mitochondrial homeostasis is regulated by the coordinated activity of different QC mechanisms that include the UPS, MDVs, and exosomes^23^. We showed that one pool of Slc25A4 is targeted for proteasomal degradation by CRL5^Ozz^, while another pool of Slc25A4 is loaded by Alix into exosomes, unveiling a cooperative action of the UPS and exosomes in regulating the same substrate, Slc25A4, in skeletal muscle. Affecting the UPS machinery by ablation of Ozz reduces Slc25A4 turnover in the tissue and shifts the QC towards Alix-mediated cargo loading into exosomes. In contrast, Alix deficiency reroutes Slc25A4 to the UPS. In both conditions, steady-state levels of Slc25A4 are affected, leading to the loss of mitochondrial proteostasis and mitochondrial dysfunction. This study has enabled us to begin to understand the interplay between UPS and exosome sorting in the regulation of mitochondrial metabolism in skeletal muscle.

Despite the great progress made towards understanding the molecular bases of mitochondrial myopathies, the etiology of many of those conditions remains unknown. Our research establishes an experimental model for future studies that address the role of Ozz and Alix in human mitochondrial myopathies, thereby providing a platform to elucidate the clinical significance of these two regulatory proteins.

## ACKNOWLEDGMENTS

We thank G. Campbell of the Light Microscopy-Cellular Imaging Shared Resource, R. Wakefield of the Electron Microscopy-Cellular Imaging Shared Resource, and R. Mosca of the Protein Production Shared Resource. A. d’Azzo holds an endowed chair in Genetics and Gene Therapy from the Jewelry Charity Fund. This study was funded in part by the National Institutes of Health grants AR049867, GM104981 and CA021764, the Assisi Foundation of Memphis, and the American Lebanese Syrian Associated Charities (ALSAC). The content is solely the responsibility of the authors and does not necessarily represent the official views of the National Institutes of Health.

## AUTHOR CONTRIBUTION

Y.C. conceived the study, designed the experiments, analyzed the data, generated the mouse models, and coordinated the efforts of all authors. G.P. designed, performed, and analyzed the soleus metabolomics data. R.R. helped with mitochondria OCR and analysis of ΔΨm data. X.Q. purified anti-Ozz and anti-Alix antibodies and helped with myoblast isolation and cultures. D.v.d.V. helped develop the procedure for isolating muscle vesicles. E.G., maintained the animal colony and performed surgeries. J.W. wrote the macros for ImageJ software to analyze MyHC content and quantify ATPase and fiber types. J.D. performed proteomic analysis of the data generated. T.B. and J.T. O. provided intellectual insights. G.C.G. provided intellectual insights and advised about the CRISPR/CAS9 mutagenesis. A. d’A. conceived the study, designed the experiments, analyzed the data, oversaw all experiments, coordinated the efforts of all authors, and secured the funding for this project.

## DECLARATION OF INTERESTS

The authors declare no competing interests.

## STAR METHODS

### Mice

Mice of all genotypes were accommodated in the Animal Resource Center at St. Jude Children’s Research Hospital. WT mice were C57Bl/6xDBA2 (B6D2F1; Jackson Labs); *Alix^−/−^*^34^ and *Ozz^−/−^*^31^ mice carry the homozygous deletion of *Alix* or *Ozz* on a C57Bl/6 background. Animals were housed in a fully AAALAC (Assessment and Accreditation of Laboratory Animal Care)-accredited animal facility. Food and water were provided ad libitum. All procedures in mice were performed in compliance with our animal protocols approved by the St. Jude Institutional Animal Care and Use Committee and per the National Institutes of Health guidelines.

### Generation of Ozz*^−/−^* mice via CRISPR/Cas9

To engineer Ozz*^−/−^* mice via CRISPR/Cas9, we synthesized single-guide RNA (sgRNA 5[-GCTGCTCCCTCCGAACACGT-3[) from linear DNA templates (IDTDNA) by using the TranscriptAid T7 High Yield Transcription Kit (Thermo Fisher Scientific). The sgRNA product was purified following the manufacturer’s instructions of TranscriptAid T7 High Yield Transcription Kit (Thermo Fisher Scientific). The sgRNA (50 ng/μL) was microinjected into the pronucleus of fertilized embryos from Rosa26-Cas9 knock-in female mice (The Jackson Laboratory), and viable embryos were implanted into oviducts of pseudo-pregnant C57BL/6J females (The Jackson Laboratory). *Ozz^−/−^*mice were born from a heterozygous intercross and used for phenotypic analyses, in parallel with age- and sex-matched WT littermates as a control group.

### Detection of genetic modification by PCR

To assess the genomic modification of *Ozz*-CRISPR/Cas9, we isolated genomic DNA from tail biopsies and used it as a template for PCR. The samples were digested in 200 μL of 50 mM NaOH and incubated at 98 °C for 1 hour. Samples were neutralized with 20 μL of 1M Tris-HCl (pH 8.0), and PCR analysis was performed with 1 μL genomic DNA in a 50-μL mixture containing 1× Green GoTaq Reaction Buffer (PROMEGA), 10 mM dNTP mix (Thermo Fisher Scientific), 50 pMol of each primer Ozz-For 5[-GTGACAAGCGAGAAAGCGGCGC-3 [and Ozz-Rev 5[-CTGCCTCCGCTTCTGGTTGACC-3 [and 1.25 U of recombinant Taq polymerase (Thermo Fisher Scientific). The PCR conditions were as follows: initial denaturation (94 °C for 3 min), 34 cycles of denaturation (94 °C for 30 s), annealing (61 °C for 30 s), extension (72 °C for 1 min), and a final extension at 72 °C for 10 min. The PCR products were run on agarose gels, purified using a gel extraction kit (QIAGEN), and sequenced to determine the location of the mutation. To discriminate between the different mutations, we subcloned the PCR products of selected sequences and amplified them in bacteria (WHS-one). DNA was extracted from single clones by using a MiniPrep Extraction Kit (QIAGEN) and Sanger-sequenced. CRISPR/Cas9 mutant mice were genotyped as follows: 8-μL PCR reactions were incubated for 2 h at 37 °C, in a mixture of 2 μL 10× Cutsmart buffer, 1 μL BsaAI enzyme (New England Biolabs), and 9 μL water. The digested products were visualized on 2% agarose gel stained with SYBR SAFE DNA gel stain (Thermo Fisher Scientific). Gels were imaged using a ChemiDoc MP Imaging System (Bio-Rad).

### Antibodies and reagents

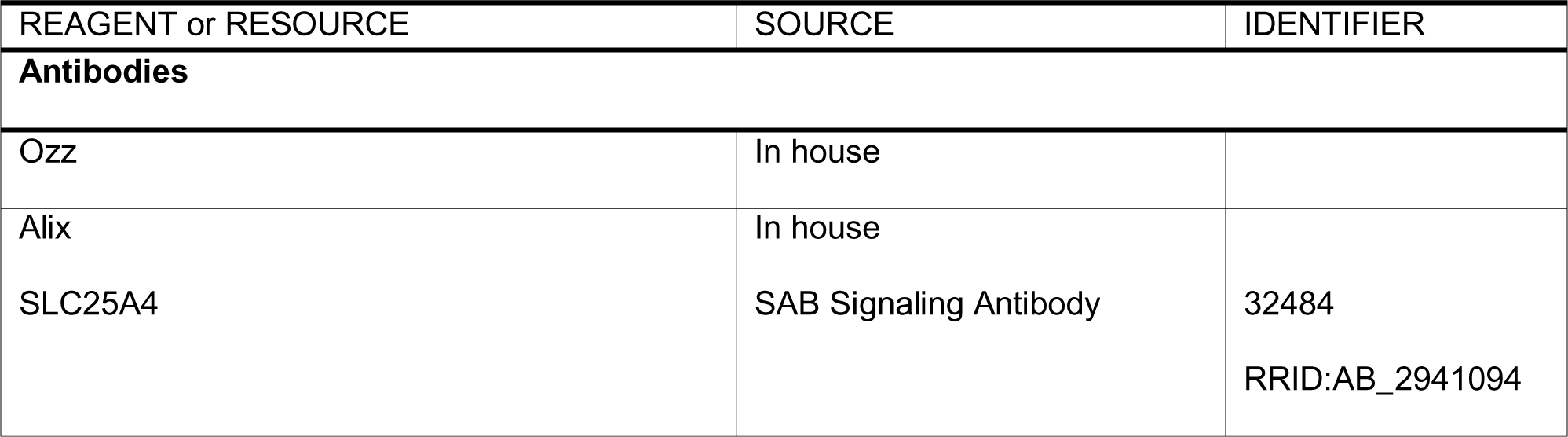
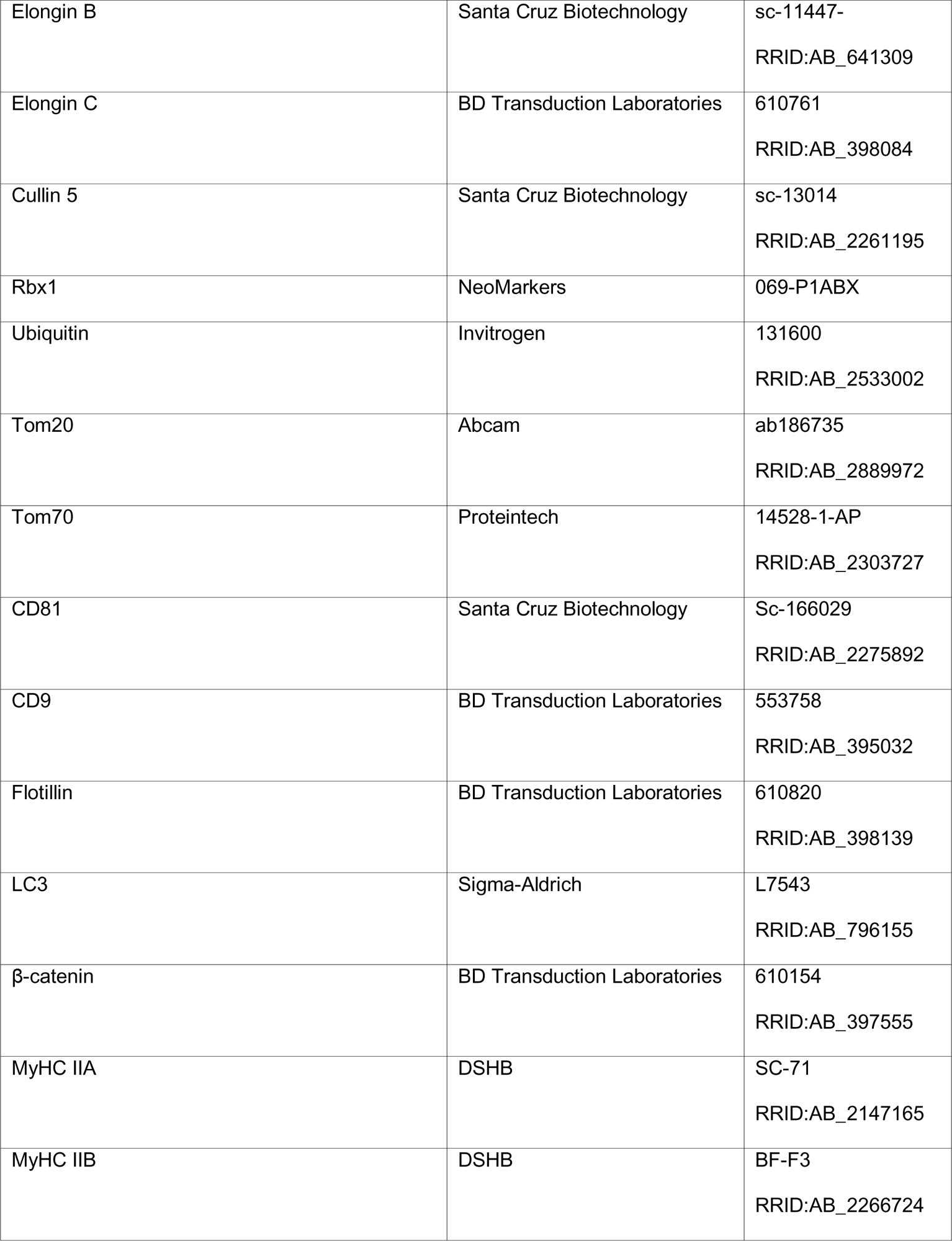
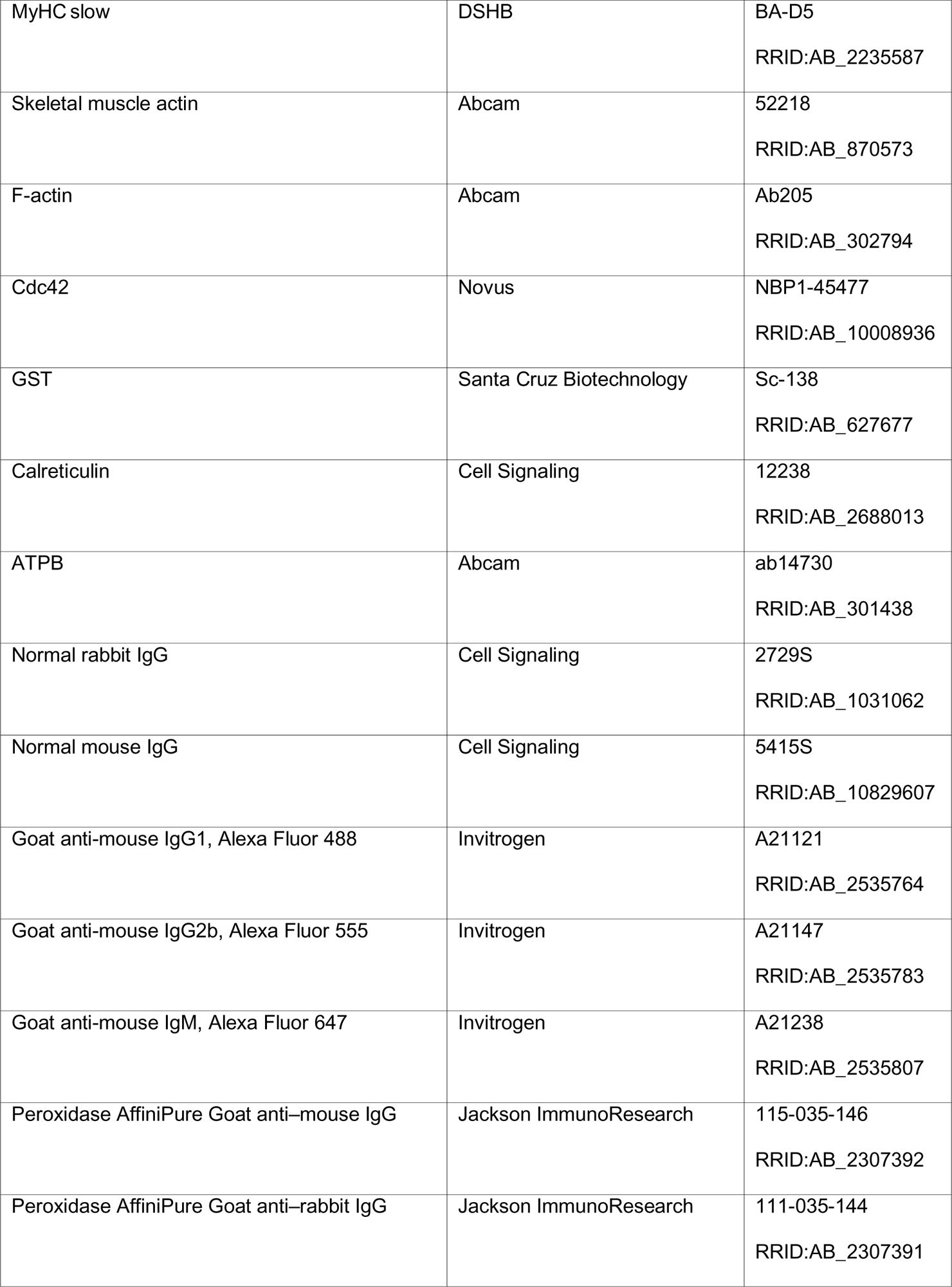
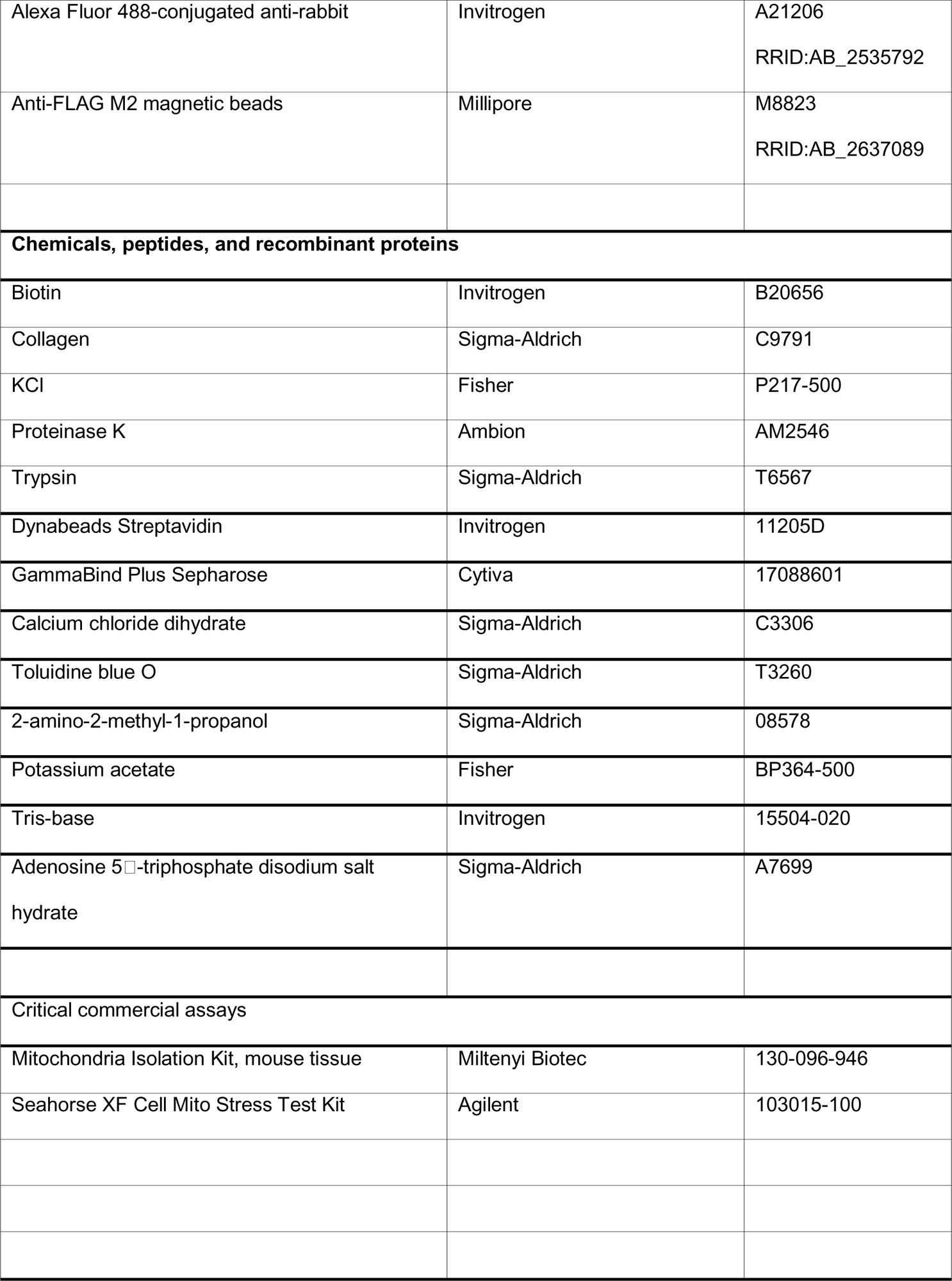
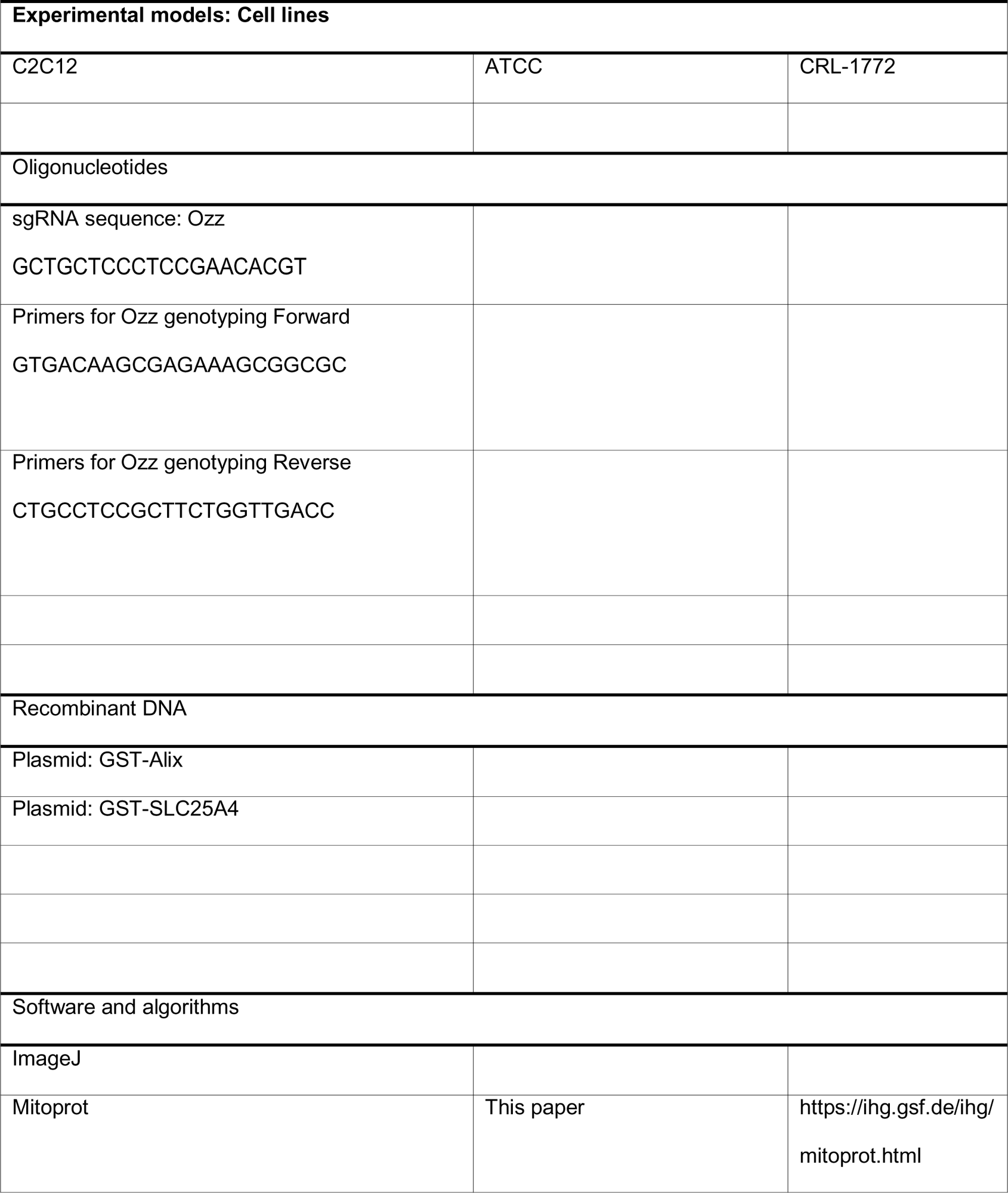

### Tissue preparation for cryosectioning

Dissected soleus or gastrocnemius muscle was embedded in 2%Tragacanth/1× PBS and flash-frozen for 10 s in isopentane cooled in liquid nitrogen. Tissues were stored at −80 °C until sectioning. Coronal or longitudinal serial sections of frozen muscles were cut on a cryostat (Leica CM3050) and adhered to superfrost plus microscope slides for immunofluorescence (8 µm) or SDH, Cyt C, and ATPase staining (12 µm). Slides were stored at –20 °C. The sections were also stained with H&E for overall morphologic assessment.

### Toluidine blue staining

Soleus dissected muscles were postfixed in 1% OsO_4_ and *en bloc* stained with 1% uranyl acetate. After standard dehydration, samples were infiltrated and embedded in Spurr low-viscosity resin (Electron Microscopy Sciences) and polymerized at 60 °C for 18 h. Semithin sections (1.5 μm) were cut and dried overnight at 45 °C. For toluidine blue staining, sections were placed on a heating tray at 95 °C, 0.1% toluidine blue was added, and the sections were incubated for 1 min. The slides were washed twice for 1 min in 95% ethanol and twice for 1 min in 100% ethanol. The sections were then cleared twice for 3 min in xylene before they were covered with paramount. Images were acquired on a Nikon C2 confocal microscope using NIS Elements software.

### Electron microscopy

Mice (aged 1–2 months) were anesthetized using Avertin (0.5 mg/g body weight) and perfused with 4% paraformaldehyde. The soleus or gastrocnemius muscles were dissected and postfixed in 2.5% glutaraldehyde in 0.1 M sodium cacodylate buffer. Both muscles were cut into small pieces, postfixed in 2% osmium tetroxide, and dehydrated via a graded series of alcohol. They were then cleared in propylene oxide, embedded in epon araldite, and polymerized overnight at 70 °C. Sections (70-nm thick) were cut on a Leica Ultracut E. The unstained sections were imaged on a JEOL 1200 EX transmission electron microscope with an AMT 2K digital camera.

### Preparation of crude or pure mitochondria from muscle tissue

Pure mitochondria were isolated per an adaptation of the protocol developed by Wieckowski et al.^66^. In brief, mice were euthanized by CO_2_ inhalation, and the soleus and gastrocnemius muscles were dissected, washed immediately in 2 mL ice-cold IB3 buffer (225 mM mannitol, 75 mM sucrose, 30 mM Tris-HCl [pH 7.4]) and stored in ice-cold IB3 buffer. Scalpels were used to cut the muscles into very small pieces, and minced samples were transferred to a 5-mL glass homogenizer tube. The samples were homogenized in 2 mL (1:5 ratio to sample) of ice-cold IB1 buffer (225mM mannitol, 75mM sucrose, 30mM Tris-HCl [pH 7.4], 0.5% BSA, 0.5mM EGTA) by using a Teflon pestle (8 strokes). The homogenates were transferred to a 5-mL Eppendorf centrifugation tube and centrifuged at 700 ×*g* for 5 min at 4 °C. The pellets were discarded, and the supernatants were centrifuged again at 700 ×*g* for 5 min at 4 °C. The resulting supernatants were then transferred to new tubes and centrifuged at 9,000 ×*g* for 10 min at 4 °C. The pellets containing crude mitochondria were gently resuspended in 2 mL ice-cold IB2 buffer (225 mM mannitol, 75 mM sucrose, 30 mM Tris-HCl [pH 7.4], 0.5% BSA) by using a Dounce homogenizer. The crude mitochondrial suspensions were further centrifuged at 10,000 ×*g* for 10 min at 4 °C, and the resulting pellets were gently resuspended in 2 mL IB3 buffer. The suspensions were centrifuged again at 10,000 ×*g* for 10 min at 4 °C, and the crude mitochondrial pellets were resuspended in 2.5 mL ice-cold mitochondrial reaction buffer (MRB; 250 mM mannitol, 5 mM HEPES [pH 7.4], 0.5 mM EGTA) by using a Dounce homogenizer. The crude mitochondrial suspensions were layered over an 8-mL Percoll solution (225 mM mannitol, 25 mM HEPES [pH 7.4], 1 mM EGTA, 30% Percoll), and topped by 1.5 mL MRB. The Percoll column was centrifuged at 95,000 ×*g* for 30 min at 4 °C. The resulting upper layer, which contained mitochondria-associated endoplasmic reticulum membranes, and the lower layer, which contained the pure mitochondrial fraction, were gently pipetted and diluted 10 times with MRB. Both fractions were then centrifuged at 6,250 ×*g* for 10 min at 4 °C. The corresponding pellets were discarded, and the supernatants containing mitochondria-associated endoplasmic reticulum membranes were transferred to ultra-clear tubes and ultracentrifuged at 100,000 ×*g* for 1 h (51-Ti rotor, Beckman) at 4 °C. The resulting pellets were snap-frozen and stored at –80 °C until use. For the mitochondrial fractions, the pellets were gently resuspended in 2 mL MRB and centrifuged at 6,250 ×*g* for 10 min at 4 °C. The resulting pellets were snap-frozen and stored at –80 °C until use.

### Sub fractionation of skeletal muscle mitochondria

Dissected gastrocnemius and soleus muscles were immediately washed with 2 mL fresh ice-cold MIB buffer (220 mM mannitol, 75 mM sucrose, 10 mM HEPES-KOH [pH 7.4], 10 mM KCl, 1 mM EDTA, 1 mM EGTA, 0.1% BSA, and protease inhibitors). Muscles were cut into small pieces using scalpels, and minced tissues were transferred to a 5-mL glass homogenizer. Ice-cold MIB buffer was added, and the muscles were homogenized using a Teflon pestle (15 strokes). The homogenates were transferred to chilled centrifuge tubes and incubated for 30 min on ice. The samples were centrifuged twice at 700 ×*g* for 5 min at 4 °C. The supernatants were transferred to new tubes and centrifuged at 5,500 ×*g* for 15 min at 4 °C. The pellets were gently resuspended in an equal volume of ice-cold MITO buffer (220 mM mannitol, 75 mM sucrose, 10 mM HEPES-KOH [pH 7.4]). A 20-µL sample of the resuspended pellet suspension was centrifuged at 5,500 ×*g* for 15 min. The resulting mitochondrial pellets were lysed in 20 μL distillated water and then snap frozen. The protein concentrations were determined by OD595 using BSA solution (Pierce). Aliquots (40 µg) of the mitochondrial preparation were placed into six new tubes with a total volume of 100 µL MITO buffer. The following solutions were then added to each tube: 2 μL of 10 mg/mL proteinase K; 700 μL of 20 mM KCl; 700 μL of 20 mM KCl + 8 μL proteinase K; 100 μL of 2% SDS + 2 μL proteinase K; 40 μL of 0.5 µg/μL trypsin. All reactions were incubated for 1 h on ice and mixed every 10 min. For the alkaline treatment, 10 μL of 1 M sodium bicarbonate was added to the sixth tube; the reaction was incubated for 30 min on ice and mixed every 10 min. After incubation, the samples were centrifuged at 15,000 ×*g* for 15 min, and the pellets were resuspended in 100 μL MITO buffer supplemented with 5 μL of 100% TCA. The samples were incubated at 60 °C for 5 min, cooled on ice for 5 min, and centrifuged at 14,000 rpm at 4 °C for 15 min. The supernatants were aspirated without disturbing the pellets. The pellets were resuspended in 18 μL of 1.5× loading buffer, supplemented with 9 μL of 0.1 M DTT. Samples were run on SDS-polyacrylamide gels and immunoblotted with selected antibodies.

### Maintenance of primary myoblast cells in culture

Myoblast cultures were established as described previously^31, 67^. In brief, primary myoblasts from WT*, Alix^−/−^*, and *ccOzz^−/−^*mice were isolated and maintained on collagen I–coated dishes (BD Biosciences) in Ham’s F-10 medium supplemented with 20% fetal calf serum (FCS), 1 ng/mL human basic fibroblast growth factor (Promega), penicillin/streptomycin (Pen/Strep; both at 100 mg/mL) (Gibco-Invitrogen) at 37 °C in 5% CO_2_. To differentiate the cells into myotubes, the culture medium was replaced with Dulbecco’s Modified Eagle Medium (DMEM) supplemented with L-glutamine and 2% horse serum.

### Mitochondrial respiration analysis

Mitochondrial respiration assays were performed using a Seahorse XF96 Cell Mito Stress Test Kit (Seahorse Biosciences), in accordance with the manufacturer’s instructions. Primary myoblasts were seeded (22,000 cells/well) in triplicates into a collagen I–coated 96-well plate in Ham’s F-12 media supplemented with 20% FCS, 1% L-glutamine, 1% Pen/Strep, and 2.5 ng/mL human fibroblast growth factor for 24 h (Day 0). Myoblasts were differentiated for 3 days in DMEM supplemented with 2% horse serum, 1% L-glutamine, and 1% Pen/Strep to form myotubes. One day before analysis, sensor cartridges were hydrated in calibration buffer and placed in a 37 °C incubator without CO_2_. On the day of the experiment, the differentiation medium was changed to Seahorse XF Base Medium (pH 7.4) containing 4 mM glutamine, 5.5 mM glucose, 2 mM pyruvate, and 5 mM HEPES. Myotubes were placed in a 37 °C incubator without CO_2_ for 1 h prior to analysis. The OCR, which represents basal respiration, was measured without any additions to the medium. Next, oligomycin was added to inhibit ATP synthase (complex V) and the OCR was recorded. The maximal OCR was measured after the uncoupler FCCP was added, and the nonmitochondrial OCR was measured by adding a mixture of rotenone/antimycin A, which shut down mitochondrial respiration. After the analysis, cells were stained with Hoechst and imaged on an XF96 extracellular flux analyzer (Seahorse Bioscience). OCR results were normalized to nuclei counts and were analyzed using Wave (software for XFp Analyzer, Agilent). Statistical calculations and graphical representations were created using GraphPad software and Microsoft Excel software.

### Measurement of mitochondrial membrane potential during myoblast differentiation

WT, *Alix^−/−^*, and *ccOzz^−/−^*primary myoblasts were seeded (5.0 × 10^4^ cells/well) in triplicates on collagen I–coated 24-well plates and maintained in culture in DMEM supplemented with 20% FCS, 1% L-glutamine, and 1% Pen/Strep at 37 °C/5% CO_2_ for 24 h (Day 0). Myoblasts were differentiated for 3 days with DMEM supplemented with 2% horse serum, 1% L-glutamine, and 1% Pen/Strep and maintained in culture at 37 °C/10% CO_2_. To measure the mitochondrial membrane potential (ΔΨm), cells were incubated at 37 °C for 30 min in media containing TMRE (150 mM). Cells were detached with 0.05% trypsin-EDTA for 5 min at 37 °C and washed twice in phenol red–free DMEM. Cellular fluorescence was detected by flow cytometry. After measuring the TMRE fluorescence, we treated the cells with FCCP (5 μM final concentration) and incubated them at RT in the dark for 5 min. The dissipation time points of the fluorescence intensity were analyzed by flow cytometry. The results obtained from the flow cytometer were analyzed using FlowJo software (TreeStar).

### Modified Gomori trichrome staining

Consecutive 12-μm sections of unfixed fresh-frozen muscle (WT, *Alix^−/−^*, and *ccOzz^−/−^*) were prepared on a cryostat, stained with Harris’ hematoxylin for 5 min, washed with water, and then stained with modified Gomori trichrome stain (0.6 g chromotrope 2R, 0.3 g Fast Green FCF, 0.6 g phosphotungstic acid, 1 mL glacial acetic acid in 100 mL deionized water). The pH was adjusted to 3.4 by using 10 N NaOH for 20 min. The slides were rinsed once in 0.2% acetic acid for 10 s, dehydrated by immersion in ethanol for 2 min (95%, 100%), and cleared twice in xylene by incubating the slides for 3 min. The cross sections were mounted with cytoseal XYL solution. Images were acquired on a Leica microscope using Leica application Suite V4.

### Succinate dehydrogenase staining

Cryosections were incubated in SDH-staining buffer (0.2 M phosphate buffer [pH 7.4], 1 mg/mL nitro-blue tetrazolium) at 37 °C for 1 h. The slides were washed thrice with deionized water. Sections were mounted in water-soluble mounting medium. Images were acquired on a Leica microscope using Leica application Suite V4.

### Cytochrome C staining

Cryosections were incubated with Cyt C–staining solution (0.2M phosphate buffer [pH 7.4], 0.5 mg/mL DAB, 1 µL/mL of 50 mM catalase solution, 1 mg/mL cyt C type II, 75 mg/mL sucrose) at 37 °C for 3 h. The slides were washed thrice with deionized water and dehydrated in ethanol (50%, 70%, 80%, [2×] 95%, [2×] 100%). The slides were cleared twice in xylene and mounted with cytoseal XYL solution. Images were acquired on a Leica microscope using Leica application Suite V4.

### Metabolic profiling analysis of the soleus-extraction of hydrophilic metabolites from skeletal tissues

Soleus muscles of 8-week-old male mice were collected, washed once in ice-cold saline, and patted dry. Next, the muscle was flash-frozen in liquid nitrogen and stored at −80 °C. To extract metabolites with different polarity, we used an adapted three-phase solvent system to obtain total hydrophilic metabolites and lipids ^68^. Briefly, the soleus muscles were weighed and then 1.2 mL chloroform/methanol/water (3:4:1, v/v/v) was added to the tissues. Approximately 25 1-mm zirconia beads were added to the samples and homogenized for 30 s at 8 m/s using a Bead Ruptor Elite (OMNI International). Next, 150 µL ice-cold saline was added, and the samples were mixed again in the Bead Ruptor. The homogenates were allowed to rest at 4 °C for 30 s and were centrifuged at 21,000 ×*g* at 4 °C for 10 min. The upper aqueous phase was transferred into new tubes, frozen on dry ice, and lyophilized. The dried extracts of hydrophilic metabolites were dissolved using 5 µL water/acetonitrile/formic acid (95:5:0.1, v/v/v) for every milligram of tissue processed, transferred into autosampler vials and 2 µL per injection was analyzed by liquid chromatography–mass spectroscopy (LC-MS). The lower organic-phase layers were transferred to new tubes and evaporated to dryness under a steam of nitrogen at RT. The dried extracts of total lipids were dissolved using 5 µL chloroform/methanol (2:1, v/v) for every milligram of tissue processed, transferred into autosampler vials, and then 10 µL per injection was analyzed by LC-MS.

### LC-MS profiling of hydrophilic metabolites

A Vanquish Horizon UHPLC (ThermoFisher Scientific) was used for the LC separations, using stepped gradient conditions as follows: 0–16.5 min, 1% to 50% B; 16.5–18 min, 50% to 99% B; 18–36 min, 99% B; 36–39 min, 99% to 1% B; 39–45 min, 1% B. Mobile phase A was water supplemented with 10 mM ammonium formate and 0.1 % formic acid. Mobile phase B was acetonitrile with 0.1% formic acid. The column used was an Acquity UPLC BEH Amide Column (1 × 150 mm, 1.7 μm) (Waters Corp.), operated at 40 °C. The flow rate was 50 μL/min, and the injection volume were 2 μL. A Thermo Scientific Q Exactive Hybrid Quadrupole-Orbitrap mass spectrometer (QE-MS) (Thermo Fisher Scientific) equipped with a HESI-II probe was employed as the detector. For each sample, two chromatographic runs were carried out to acquire separate data for negative ions and positive ions. The QE-MS was operated using a data-dependent LC-MS/MS method (Top10 dd-MS2) for both positive and negative ion modes. The MS was operated at a resolution of 140,000 (FWHM, at m/z 200), AGC targeted of 3 × 10^6^, max injection time of 100 ms. The instrument’s operating conditions were scan range 60-900 m/z, sheath gas flow 20, aux gas flow 5, sweep gas 1, spray voltage 3.6 kV for positive mode and 2.5 kV for negative mode, capillary temp 320 °C, S-lenses RF level 55, aux gas heater 320 °C. For the Top10 dd-MS2 conditions, a resolution of 35,000 was used, AGC targeted of 1 × 10^5^, max injection time 60 ms, MS2 isolation width 1.0 m/z, NCE 35.

### Data processing, metabolite identification, and statistics

The software Compound Discoverer 3.1 (CD3.1) (Thermo Fisher Scientific), XCMS Online, and MetaboAnalyst 5.0 were used to process untargeted metabolomic analyses. For CD3.1, a predefined workflow from the software was employed: Untargeted metabolomics with statistics detects unknowns IDs by using online databases and maps compounds to biological pathways by using Metabolika software. The spectra-alignment node used the adaptative curve algorithm, with a maximum shift of 2 min. The compound-detection option was set for a signal-to-noise ratio ≥3, minimum peak intensity 1 × 10^6^, parent ion mass tolerance of ±5 ppm, RT tolerance of 0.25 min, and preferred fragments data selection M+H, M-H. The data were normalized by the constant mean algorithm. The metabolite identification and pathway analysis nodes used the default features in this metabolomics workflow. Complete metabolomics workflows for multigroup and pairwise jobs were run using XCMS Online. For positive-mode data files, the UPLC/Q-Exactive (3110) parameter was used, and for negative-mode data files, a custom set of parameters was created based on param. #3110 to identify adducts in the negative-mode files. The XCMS Pathway Cloud Plot’s p-value threshold filter was set to 0.05, and then the overlapping metabolites from all filtered metabolic pathways were tabulated (positive and negative modes combined). They were then used to perform metabolite set enrichment and metabolic pathways analysis in MetaboAnalyst 5.0.

Data represent the means ± standard error of the mean. The differences between treated groups and untreated groups were analyzed using CD3.1, LipidSearch v4.1, and MetaboAnalyst v5.0. Statistical differences were considered significant for *p-*values ≤0.05; NS denotes nonsignificant differences.

### Immunofluorescence and imaging

Cross sections of soleus muscle were fixed and permeabilized with 1% PFA, 0.4% Triton-X 100 in 1× PBS for 15 min at RT. Sections were blocked for 1 h in blocking buffer (PBS containing 5% normal donkey serum, 0.1% BSA, and 0.4% Triton X-100) and then incubated with primary antibodies (Table) diluted in blocking buffer at 4 °C overnight. Next day, the slides were washed thrice with washing buffer (PBS containing 0.4% Triton X-100) and incubated with secondary antibody diluted in blocking buffer at 4 °C for 1 h. Following three washings, the sections were mounted with ProLong Gold antifade reagent with DAPI. Images were acquired on a Nikon C2 confocal microscope using NIS Elements software. A macro was developed in ImageJ to count the different types of myosin fibers (Type I, IIa, IIb) in the soleus. A threshold for detection was set to quantify the distinct fibers per field and counts from all images using the same parameters were batched.

### Determination of ATPase activity

Frozen serial sections of the soleus were air dried at RT for 5 min. Slides were incubated in acid preincubation buffer (50 mM potassium acetate, 17.7 mM CaCl_2_ dihydrate [pH 4.5], with glacial acetic acid and filtered at 0.22 µm) at RT for 2 min and washed thrice with Tris-buffer (100 mM Tris base, 17.7 mM CaCl_2_ dihydrate [pH 7.8], with 5 N HCl and filtered at 0.22 µm) for 2 min. The slides were transferred to a container with basic incubation buffer (20 mM KCl, 2.7 mM ATP, disodium salt, 50 mM CaCl_2_ dihydrate, 100 mM 2-amino-2-methyl-propanol HCl [pH 10.5], with 1 N HCl, not filtered) and incubated at 24 °C for 25 min. The slides were rinsed four times in three changes of 1% CaCl_2_ and stained in 0.1% toluidine blue for 45 s. The slides were rinsed in distilled H_2_O for 30 s and dehydrated by dipping five times in serial concentrations of ethanol (95%, 100%, 100%). The slides were cleared twice in xylene for 3 min and mounted with cytoseal XYL. Images were acquired on a AxioScan Z.1 Whole Slide Scanner using Zen Black (Zeiss) software. A macro was developed in ImageJ to count the different types of myosin fibers (Type I, IIa, IIb) in the soleus. A threshold for detection was set to quantify the distinct fibers per field and counts from all images using the same parameters were batched.

### Generation, transduction, and purification of biotinylated Ozz

The biotinylating procedure was performed as described previously^53, 54^. *Ozz* cDNA was cloned in the pBABE-puro vector containing the 23–amino acid biotinylation tag (Avi-tag). Retrovirus particles were produced by transfecting the packaging cell line Eco Phoenix with Avi-pBABE-puro or Avi-Ozz-pBABE-puro plasmids using the transfection reagent FuGENE 6. BirA-C2C12 cells, generously provided by G. Grosveld of St. Jude Children’s Research Hospital, were transduced with Avi-pBABE-puro or Avi-Ozz-pBABE-puro retroviruses for 48 h, followed by neomycin (0.8 mg/mL) and puromycin (3 μg/mL) selection for 5 days. Selected cells were maintained in DMEM plus medium (DMEM supplemented with 20% FCS, L-glutamine, 10^4^ U/mL/10^4^ _μ_g/mL penicillin/streptomycin, 0.8 mg/mL neomycin, and 1.0 μg/mL puromycin). For purification of biotinylated proteins, 1.0 × 10^6^ C2C12 cells were seeded in 150 cm^2^ flasks and grown in DMEM plus medium containing 50mM of biotin (Day 0). For differentiated C2C12 cells, 3.0 × 10^6^ C2C12 cells were seeded in 150 cm^2^ flasks and grown in DMEM plus medium, which was changed after 48 h for DMEM medium supplemented with 2% horse serum, L-glutamine, 10^4^ U/mL/10^4^ _μ_g/mL penicillin/streptomycin, 0.8 mg/mL neomycin, 1.0 μg/mL puromycin, and 50 mM of biotin and incubated for 72 h. (Day 3). All cell cultures were grown at 37 °C in 5% CO_2_. Cell pellets were collected from Avi-Ozz–overexpressing C2C12 cells at Day 0 and Day 3. For crude mitochondria preparations, the same overexpressing cells were first lysed in ice-cold lysis buffer (1× TBS [pH 8.0], 0.3% NP-40, 20% glycerol, 0.215 mM EDTA [pH 8.0], 150 mM KCl, CelLytic Express, protease inhibitors, and phosphatase inhibitors) by using a Dounce homogenizer (15 strokes). The homogenates were kept on ice with slight agitation for 30 min and centrifuged at 16,000 ×*g* for 10 min at 4 °C. The resulting supernatants were transferred into fresh tubes, and protein concentrations were measured. Paramagnetic streptavidin beads were blocked by washing them thrice in 1× TBS with 200 ng/μL purified chicken serum albumin (Sigma-Aldrich). Approximately 20 μL of beads were added per 1 mg of soluble extract. Binding of the samples to the beads was done in a solution of 1× TBS (pH 8.0), 0.3% NP-40, 20% glycerol, 0.215 mM EDTA (pH 8.0), 150mM KCl, at 4 °C under agitation from 1 h to overnight. The samples were washed six times in binding solution at RT. Bound proteins were released by boiling the beads in sample buffer and then identified by SDS-PAGE under denaturing conditions and immunoblotting with selected antibodies.

### Generation, transduction, and purification of FLAG-Alix

C2C12 cells were transduced with MSCV-GFP empty vector or MSCV-GFP expressing N-terminal FLAG-tagged full-length Alix. At 72 h after transduction, cells were sorted and selected by GFP expression. A total of 1.0 × 10^6^ C2C12 cells were seeded into 150 cm^2^ flasks. Proliferating cells were grown in DMEM supplemented with 20% FCS, L-glutamine, 10^4^ U/mL/10^4^ _μ_g/mL penicillin/streptomycin and differentiated C2C12 cells were grown in DMEM supplemented with 2% horse serum, L-glutamine, and 10^4^ U/mL//10^4^ _μ_g/mL penicillin/streptomycin. Cells were grown at 37 °C in 5% CO_2_.

Mitochondrial preparations from C2C12 cells overexpressing FLAG-tagged full-length Alix were lysed with ice-cold lysis buffer (1× TBS [pH 8.0], 0.3% NP-40, 20% glycerol, 0.215 mM EDTA [pH 8.0], 150 mM KCl, protease inhibitors, and phosphatase inhibitors) by using a Dounce homogenizer (15 strokes). The homogenates were kept on ice with slight agitation for 30 min and centrifuged at 20,000 rpm for 10 min. The resulting supernatants were transferred into clean tubes and protein concentration was measured. Samples were applied to pre-equilibrated anti-FLAG M2 affinity gel and incubated at 4 °C for 1 h. The incubated resin was collected by centrifugation at 1,500 rpm for 5 min at 4 °C and washed with lysis buffer. After extensive washing, the FLAG-tagged protein was eluted using 3× FLAG peptide (St. Jude Peptide Synthesis Facility), and eluted fractions were subjected to SDS–PAGE under denaturing conditions followed by immunoblotting with the indicated antibodies.

### Coimmunoprecipitations

For immunoprecipitation experiments, crude mitochondria preparations were lysed with 1× TEVP buffer ^69^ (10 mM Tris base, 5 mM NaF, 1 mM Na_3_VO_4_, 1 mM EDTA, 1 mM EGTA [pH 7.4]) supplemented with 0.5% Triton X-100 and protease inhibitors. Mitochondrial lysates (140 μg) were incubated with anti-SLC25A4 antibody (5 μg) at 4 °C overnight. Samples were immunoprecipitated with 40 μL Pure Proteome Protein A/G Mix Magnetic Beads coupled to V5 peptide at 4 °C for 1 h. The beads were washed thrice with ice-cold lysis buffer and once with 1× TEVP buffer. Bound proteins were released by boiling the beads in sample buffer. They were then run on SDS-polyacrylamide gels, under denaturing conditions, followed by immunoblotting with the indicated antibodies.

### In vitro ubiquitination of mitochondria proteins

The ubiquitination assay was performed by incubating 3.0 µg of a bacterially expressed GST-SLC25A4 with 150 ng purified recombinant E1, 200 ng purified recombinant E2, 1.0 µg CRL5^Ozz^ ubiquitin ligase, and 7.5 µg ubiquitin in a final volume of 30 µL ubiquitination buffer (0.05 M Tris-HCl [pH 7.6]; 0.01 M MgCl_2_; 0.004 M ATP) at 30 °C for 1 h.

To analyze the ubiquitinated products, we diluted the ubiquitination reaction in 500 µL RIPA buffer (50 mM Tris HCl [pH 7.5], 150 mM NaCl, 1% NP-40, 0.1% deoxycholate, 0.1% SDS, 1 mM EDTA, protease inhibitors, and phosphatase inhibitors) and incubated with 5 µg anti-SLC25A4 antibody at 4 °C for 1 h. Samples were immunoprecipitated with 40 µL preclear GammaBind plus sepharose at 4 °C for 1 h. The beads were washed thrice with RIPA buffer and once with resuspension buffer without detergent. Bound proteins were released by boiling the beads in sample buffer, and they were resolved on 7.5% SDS-PAGE| and immunoblotted with anti-GST or anti-ubiquitin antibodies.

### Exosome isolation from skeletal muscle

Exosomes were isolated as described previously^55^. Briefly, gastrocnemius or soleus muscles of 8-week-old mice were collected and washed once with ice-cold 1× PBS. Tissues were gently sliced into small fragments and transferred to glass homogenization tubes containing 4 mL ice-cold 1× PBS. The samples were hand homogenized at 8,000 rpm (20 strokes) and transferred to 5-mL centrifuge tubes. Exosomes were purified by sequential centrifugation steps at 300 ×*g* for 10 min, 2000 ×*g* for 10 min, and 10,000 ×*g* for 30 min to remove cells and cell debris. The supernatants were then ultracentrifuged at 100,000 ×*g* for 120 min at 4 °C. The supernatant was discarded, and the exosome pellets were washed once with cold PBS and then ultracentrifuged at 100,000 ×*g* at 4 °C for 2 h. Exosomal pellets were resuspended in 0.25 M sucrose and loaded onto a step gradient of 2 to 0.5 M sucrose in 10 mM tris-HCl buffer (pH 7.4) and 1 mM Mg (Ac)_2_. The gradients were ultracentrifuged at 100,000 ×*g* for 2.5 h in a Beckman Coulter SW41Ti rotor. Fractions were collected and precipitated with 100% TCA (110 µL). Pellets were resuspended in NuPAGE lithium dodecyl sulfate sample buffer (Life Technologies or Bio-Rad) with or without DTT and used for Western blot analyses.

### Mito-QC mouse model

The mito-QC mouse model was generated as previously described by McWilliams et al., 2016^48^.

### Statistics

Data were expressed as mean ± standard deviation^65^ and evaluated using Student’s *t*-test or Ordinary one-way ANOVA test for comparison with WT samples. Mean differences were considered statistically significant when P-values were less than 0.05.

## Notes

### Competing Interest Statement

The authors have declared no competing interest.

