## Supplemental Text for "Mitochondrial proteostasis mediated by CRL5^Ozz^ and Alix maintains skeletal muscle function"

**SUPPLEMENTAL INFORMATION**

**TABLES**

**Table S1 WT vs *Alix*^-/-^**

**Metabolite Name**

AMP

(2E,8Z,11Z,14Z,17Z)-icosa-2,8,11,14,17-pentaenoyl-CoA

(5Z,8Z,11Z,14Z,17Z)-icosapentaenoyl-CoA

Mevalonic acid

(S)-2-amino-6-oxohexanoate

2-aminoprop-2-enoate

2-iminopropanoate

2-keto-6-aminocaproate

3-acetamidopropanal

Ureidoisobutyric acid

3-ureidopropionate

4-acetamidobutanoate

4-imidazoleacetate

5,6-dihydrothymine

5,6-dihydrouracil

5-Methoxyindoleacetate

acetol

adenine

adenosine 3',5'-bisphosphate

coenzyme A

dGDP

dGMP

glycerol

histamine

IMP

inosine

lactaldehyde

lactate

L-arginine

Argininosuccinic acid

laurate

linoleate

L-saccharopine

malonate semialdehyde

melatonin

methylglyoxal

Acetyl-N-formyl-5-methoxykynurenamine

N-Acetylmannosamine

N-Acetylneuraminic acid

N-Acetylneuraminic acid 9-phosphate

nicotinamide

palmitate

pyruvate

stearate

thymine

uracil

AMP

(R)-3-hydroxybutanoate

(S)-3-methyl-2-oxopentanoate

(S)-dihydroorotate

Glyceric acid 1,3-biphosphate

1D-myo-inositol 4-monophosphate

2-aminoprop-2-enoate

2'-deoxyguanosine

2'-deoxyinosine

2-iminopropanoate

3-hydroxyanthranilate

Ureidoisobutyric acid

3-ureidopropionate

4-imidazoleacetate

4-methyl-2-oxopentanoate

5,6-dihydrothymine

5,6-dihydrouracil

acetyl-CoA

adenosine

ADP-D-ribose

glucose

D-glyceraldehyde

dGMP

1D-Myo-inositol 1,4-bisphosphate

dTMP

dTTP

dUMP

dUTP

ethanol

formate

fructose

fructose 6-phosphate

fructose 1,6-bisphosphate

fructose 2,6-bisphosphate

fumarate

galactose

galactose 1-phosphate

GDP-glucose

GDP-D-mannose

glucose 1-phosphate

glucose 6-phosphate

glycine

glyoxylate

hydrogen cyanide

imidazole acetaldehyde

IMP

keto-D-fructose

lactate

L-arginine

Argininosuccinic acid

L-citrulline

lipoate

malate

mannose 1-phosphate

mannose 6-phosphate

methylglyoxal

myo-inositol

N-Acetyl-glucosamine 1-phosphate

NAD+

N-carbamoyl-L-aspartate

octanoate

orotidine 5'-phosphate

palmitate

pyruvate

stearate

succinate

sucrose

thiosulfate

thymine

UDP

Uridine diphosphategalactose

Uridine diphosphate glucose

uracil

Trehalose

**Table S2 WT vs *Ozz^-/-^***

**Metabolite Name**

β-estradiol

19-hydroxyandrostenedione

19-oxo-testosterone

2-aminoprop-2-enoate

2'-deoxycytidine

2'-deoxyuridine

2-iminopropanoate

2-phospho-D-glycerate

3-phospho-D-glycerate

Ureidoisobutyric acid

3-ureidopropionate

4-imidazoleacetate

5,6-dihydrothymine

5,6-dihydrouracil

5,6-dihydroxyindole-2-carboxylate

acetoacetate

CDP-choline

fumarate

imidazole acetaldehyde

L-arginine

L-citrulline

L-dopachrome

L-gulonate

L-leucine

L-xylulose

methylglyoxal

palmitate

phosphocholine

phosphoenolpyruvate

pyruvate

stearate

thymidine

thymine

uracil

urea

Galactose 1-phosphate

Glucose 1-phosphate

Mannose 1-phosphate

Mannose 6-phosphate

fructose 6-phosphate

Glucose 6-phosphate

2-aminoprop-2-enoate

2'-deoxyguanosine

2'-deoxyinosine

2'-deoxyuridine

Deoxyribose 1-phosphate

2-iminopropanoate

2-oxo-4-methylthiobutanoate

2-oxoglutaramate

2-oxoglutarate

2-phospho-D-glycerate

3-phospho-D-glycerate

Ureidoisobutyric acid

3-ureidopropionate

4-Acetamidobutanoic acid

4-acetamidobutanoate

5,6-dihydrothymine

5,6-dihydrouracil

5-methylthioribulose 1-phosphate

7α,12α-dihydroxycholest-4-en-3-one

adenosine

adenosine 3',5'-bisphosphate

AMP

cis-aconitate

coenzyme A

dGDP

D-glyceraldehyde

D-glyceraldehyde 3-phosphate

dGMP

dGTP

dihydroxyacetone phosphate

ethanol

fructose 1,6-bisphosphate

fumarate

glutathione

lactaldehyde

lactate

Argininosuccinic acid

L-ascorbate

L-citrulline

Dehydroascorbic acid

linoleate

malate

octanoate

oxaloacetate

palmitate

phosphoenolpyruvate

pyruvate

5-Methylthioribulose 1-phosphate

sn-glycerol 3-phosphate

thymidine

thymine

UDP

uracil

UTP

Glucose 1-phosphate

Glucose 6-phosphate

Mannose 6-phosphate

fructose 1-phosphate

fructose 6-phosphate

Fructose 2,6-bisphosphate

**Table S3: Common metabolites in *Ozz* ^-/-^ and *Alix*^-/-^ compared to WT soleus muscle**

| 1 | 3-Phosphoglyceric acid | PGA | Glycolysis |  |
| --- | --- | --- | --- | --- |
| 2 | Acetyl-CoA | AcetylCoA |  | MCF-TCA |
| 3 | adenosine 3',5'-bisphosphate | ADP | SLC25A42 | adenosine bisphosphate |
| 4 | cis-Aconitic acid | Aconitate | Citric acid cycle | MCF-TCA |
| 5 | Citric acid | Citrate | SLC25A1 | MCF-TCA |
| 6 | Coenzyme A | CoA | SLC25A42 | MCF-TCA |
| 7 | Deoxycytidine | dC | dUMP--dTMP |  |
| 8 | Deoxyribose 1-phosphate | dR1P |  |  |
| 9 | Deoxyuridine | dU |  |  |
| 10 | Deoxyuridine triphosphate | dUTP |  |  |
| 11 | D-Fructose | Fructose |  |  |
| 12 | D-Glyceraldehyde 3-phosphate | G3P |  |  |
| 13 | Dihydrothymine | dhT | Thymine | MCF |
| 14 | Dihydrouracil | dhU | Uracil | MCF |
| 15 | Dihydroxyacetone phosphate | DHAP | Glycolysis |  |
| 16 | dUMP | dUMP | SLC25A33/36 |  |
| 17 | Ethanol | Ethanol |  |  |
| 18 | Fructose 1,6-bisphosphate | FBP |  |  |
| 19 | Fructose 6-phosphate | F6P |  |  |
| 20 | Fumaric acid | Fumarate | Intermedia in the citric acid cycle to produce ATP | TCA |
| 21 | Glucose 1-phosphate | G1P | Glycolysis |  |
| 22 | Glucose 6-phosphate | G6P | Glycolysis |  |
| 23 | Glyceric acid 1,3-biphosphate | ‎1,3BPG | Glycolysis |  |
| 24 | Guanosine diphosphate | GDP | Glycolysis |  |
| 25 | L-Dihydroorotic acid | Dihydroorotic | Pyrimidine biosynth | OXPHOS |
| 26 | L-Malic acid | Malate | SLC25A10 |  |
| 27 | Oxalacetic acid | Oxa | Glucogenesis, Urea, aa | MCF-TCA |
| 28 | Oxoglutaric acid(alpha-ketoglutarate) | Oxo | Krebs Cycle |  |
| 29 | Phosphoenolpyruvic acid | PEP | Glycolysis/Gluconeoge | Citric acid cycle-in the presence of O2, no O2 produce lactate |
| 30 | Pyruvic acid | Pyruvate | Glycolysis/Gluconeoge | Citric acid cycle-in the presence of O2, no O2 produce lactate |
| 31 | Sucrose | Sucrose |  |  |
| 32 | Thymidine | Thymidine | SLC25A33 | MCF |
| 33 | Thymidine 5'-triphosphate | TTP | SLC25A33 | MCF |
| 34 | Thymine | Thymine | SLC25A33 | MCF |
| 35 | Uracil | Uracil | SLC25A33 | MCF |
| 36 | Ureidoisobutyric acid | Ureidoisobutyric | Thymine | MCF |
| 37 | Ureidopropionic acid | Ureidopropionic | Uracyl | MCF |
| 38 | Ureidosuccinic acid | Ureidosuccinic | Pyrimidine | MCF |
| 39 | Uridine 5'-diphosphate | UDP | Glycogenesis |  |
| 40 | Uridine diphosphate glucose | UDP-Glc | Glycogenesis | A low level of UDP-Glc occurs in cells exposed to hypoxia or glucose starvation. |
| 41 | Uridine triphosphate | UTP |  | Equal to ATP |

**Table S4 Pathway analysis between WT vs *Alix*^-/-^**

| **S.no** | **Pathway Name** | **Match Status** | ***p* value** | **FDR** |
| --- | --- | --- | --- | --- |
|  | Pyrimidine Metabolism | 14/54 | 2.686E-4 | 0.015999 |
|  | Galactose Metabolism | 10/31 | 3.232E-4 | 0.015999 |
|  | Amino Sugar Metabolism | 9/31 | 0.0015536 | 0.047101 |
|  | Starch and Sucrose Metabolism | 8/26 | 0.0019031 | 0.047101 |
|  | Lactose Synthesis | 5/14 | 0.0071479 | 0.13758 |
|  | Glycolysis | 6/20 | 0.008338 | 0.13758 |
|  | Nucleotide Sugars Metabolism | 5/16 | 0.013355 | 0.18888 |
|  | Gluconeogenesis | 7/30 | 0.018874 | 0.23357 |
|  | Ketone Body Metabolism | 4/12 | 0.021249 | 0.23374 |
|  | Citric Acid Cycle | 6/25 | 0.025689 | 0.25432 |
|  | Purine Metabolism | 11/63 | 0.030554 | 0.25473 |
|  | Beta-Alanine Metabolism | 6/26 | 0.030876 | 0.25473 |
|  | Aspartate Metabolism | 7/34 | 0.036276 | 0.27626 |
|  | Fructose and Mannose Degradation | 6/28 | 0.043207 | 0.30554 |
|  | Pyruvate Metabolism | 6/30 | 0.058269 | 0.37198 |
|  | Urea Cycle | 5/23 | 0.060118 | 0.37198 |
|  | Transfer of Acetyl Groups into Mitochondria | 4/18 | 0.084256 | 0.49067 |
|  | Glutamate Metabolism | 7/45 | 0.12871 | 0.70305 |
|  | Pyruvaldehyde Degradation | 2/7 | 0.13818 | 0.70305 |
|  | Alanine Metabolism | 3/14 | 0.14203 | 0.70305 |

**Table S5 Pathway analysis between WT vs *Ozz*^-/-^**

| **S.no** | **Pathway Name** | **Match Status** | ***p* value** | **FDR** |
| --- | --- | --- | --- | --- |
|  | Gluconeogenesis | 12/30 | 2.67E-07 | 0.000026394 |
|  | Glycolysis | 9/20 | 3.04E-06 | 0.00015061 |
|  | Urea Cycle | 8/23 | 1.07E-04 | 0.0035371 |
|  | Pyrimidine Metabolism | 12/54 | 2.55E-04 | 0.0063047 |
|  | Fructose and Mannose Degradation | 8/28 | 5.04E-04 | 0.009979 |
|  | Citric Acid Cycle | 7/25 | 0.001333 | 0.022001 |
|  | Pyruvate Metabolism | 7/30 | 0.004198 | 0.059367 |
|  | Starch and Sucrose Metabolism | 6/26 | 0.008559 | 0.093747 |
|  | Pentose Phosphate Pathway | 6/27 | 0.010379 | 0.093747 |
|  | Pyruvaldehyde Degradation | 3/7 | 0.010416 | 0.093747 |
|  | Malate-Aspartate Shuttle | 3/7 | 0.010416 | 0.093747 |
|  | Lactose Synthesis | 4/14 | 0.014689 | 0.12035 |
|  | Glycerol Phosphate Shuttle | 3/8 | 0.015803 | 0.12035 |
|  | Mitochondrial Electron Transport Chain | 4/15 | 0.018945 | 0.13058 |
|  | Amino Sugar Metabolism | 6/31 | 0.020443 | 0.13058 |
|  | Glycerolipid Metabolism | 5/23 | 0.021104 | 0.13058 |
|  | De Novo Triacylglycerol Biosynthesis | 3/9 | 0.022482 | 0.13092 |
|  | Nucleotide Sugars Metabolism | 4/16 | 0.023895 | 0.13142 |
|  | Aspartate Metabolism | 6/34 | 0.031365 | 0.16343 |
|  | Beta-Alanine Metabolism | 5/26 | 0.034767 | 0.16954 |
|  | Transfer of Acetyl Groups into Mitochondria | 4/18 | 0.035964 | 0.16954 |
|  | Tyrosine Metabolism | 8/55 | 0.03955 | 0.17106 |
|  | Cardiolipin Biosynthesis | 3/11 | 0.039741 | 0.17106 |
|  | Arginine and Proline Metabolism | 7/48 | 0.052543 | 0.21674 |
|  | Galactose Metabolism | 5/31 | 0.06764 | 0.26785 |
|  | Alanine Metabolism | 3/14 | 0.07495 | 0.28539 |
|  | Purine Metabolism | 8/63 | 0.078406 | 0.28749 |
|  | Glutamate Metabolism | 6/45 | 0.10072 | 0.000026394 |

**SUPPLEMENTAL LEGENDS**

**Figure S1. Western blot analyses of different skeletal muscles**

(A) Immunoblot analyses of soleus, tibialis, gastrocnemius, and diaphragm muscles. Lysates from WT mice were probed with anti-Ozz and anti-Alix antibodies that showed the presence of both proteins in all muscles. Anti–skeletal muscle actin (SkM actin) was used as a control.

(B) Immunofluorescence staining of gastrocnemius/soleus muscle sections from WT and *Alix^−/−^* mice with antibodies specific for MyHC I and Alix. Scale bar: 50 μm

(C) Immunofluorescence staining of soleus muscle sections from WT, *Alix^−/−^* and *Ozz^−/−^* mice with antibodies specific for Ozz and Alix. Scale bar: 50 μm

(D) H&E-stained cross sections of soleus muscles from 2-month-old WT, *Alix^−/−^*, and *Ozz^−/−^* mice shows the presence of small fibers (arrowheads) in *Alix^−/−^* and *Ozz^−/−^* mouse models compared to WT. Scale bar: 50 μm.

**Figure S2. Oxygen consumption rate of primary myoblasts (Day 0)**

(A) Oxygen consumption rate (OCR) profile of WT, *Alix^−/−^*, and *ccOzz^−/−^* mouse primary myoblasts (Day 0) in response to treatment with oligomycin (ATP-linked respiration and proton leak), FCCP (mitochondrial reserve capacity), and rotenone/antimycin A (nonmitochondrial respiration). Hoechst 33342 staining was added to count the number of cells.

(B) Analysis of the different stages of myoblast respiration determined from the OCR analysis (panel A) of primary myoblasts of WT, *Alix^−/−^*, and *ccOzz^−/−^* mice (Day 0). Data presented are the mean ± SD, Student’s (unpaired) *t*-test; n = 3 independent samples.

(C) Quantitative analysis of ΔΨΜ by using the mean fluorescence intensity (MFI) of TMRE in primary myoblasts of WT, *Alix^−/−^*, and *ccOzz^−/−^* muscle. Data presented are the mean ± SD, Student’s (unpaired) *t*-test; n = 100 cells per n = 3 independent samples.

**Figure S3.** **Mitophagy analysis using mito-QC reporter mice**

Representative confocal images of soleus muscles isolated from MitoQC/WT, MitoQC/*Alix^−/−^* and MitoQC/cc*Ozz^−/−^* mice show increased mitophagy (red) in both MitoQC/*Alix^−/−^* and MitoQC/cc*Ozz^−/−^* mice. Scale bars: 10 μm.

**Figure S4.** **Exosomes contain Slc25A4**

Immunoblot analyses of WT, *Alix^−/−^*, and *ccOzz^−/−^* gastrocnemius/soleus muscle exosomes subjected to sucrose density gradient analysis and probed with anti-Ozz, anti-Alix, anti-Slc25A4, anti-CD81, and anti-F-actin antibodies.
