## Supplementary figures and images for "Mitochondrial proteostasis mediated by CRL5^Ozz^ and Alix maintains skeletal muscle function"

### Supplemental Figure 1

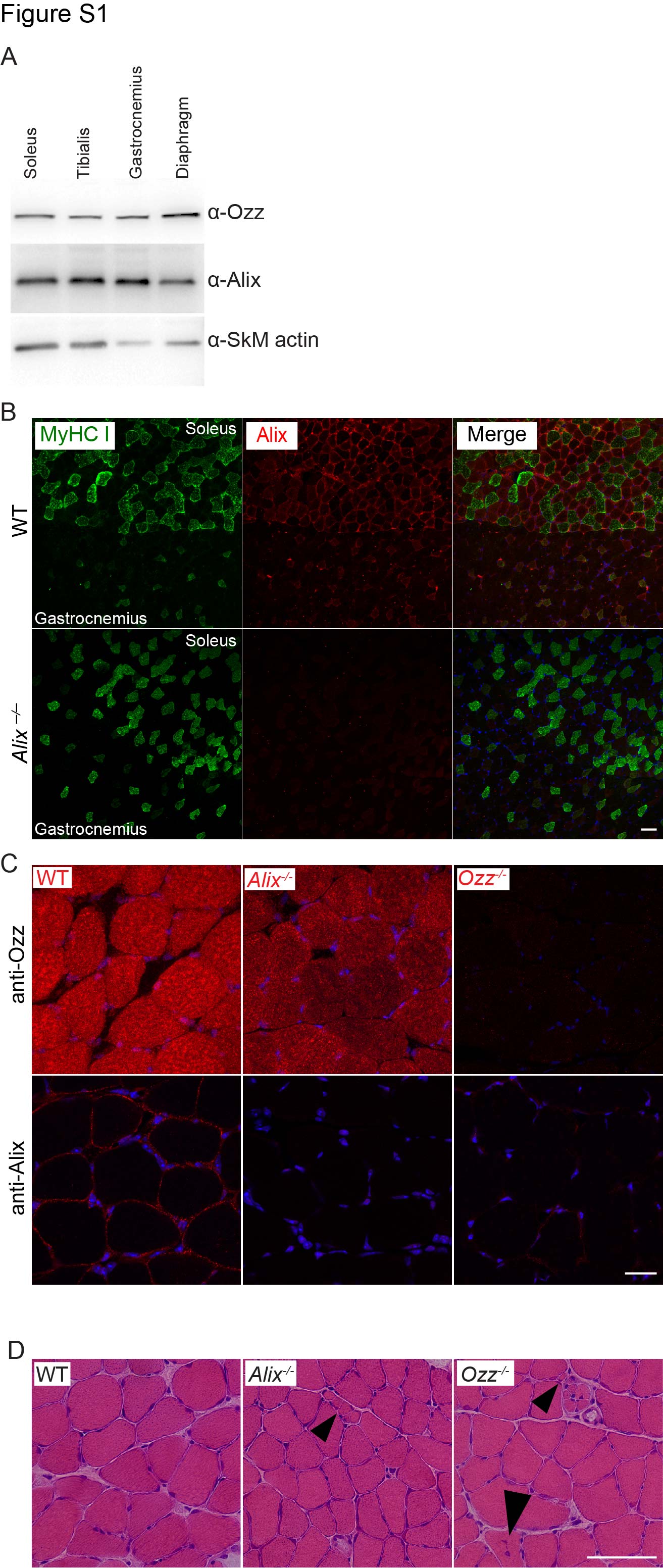

### Supplemental Figure 2

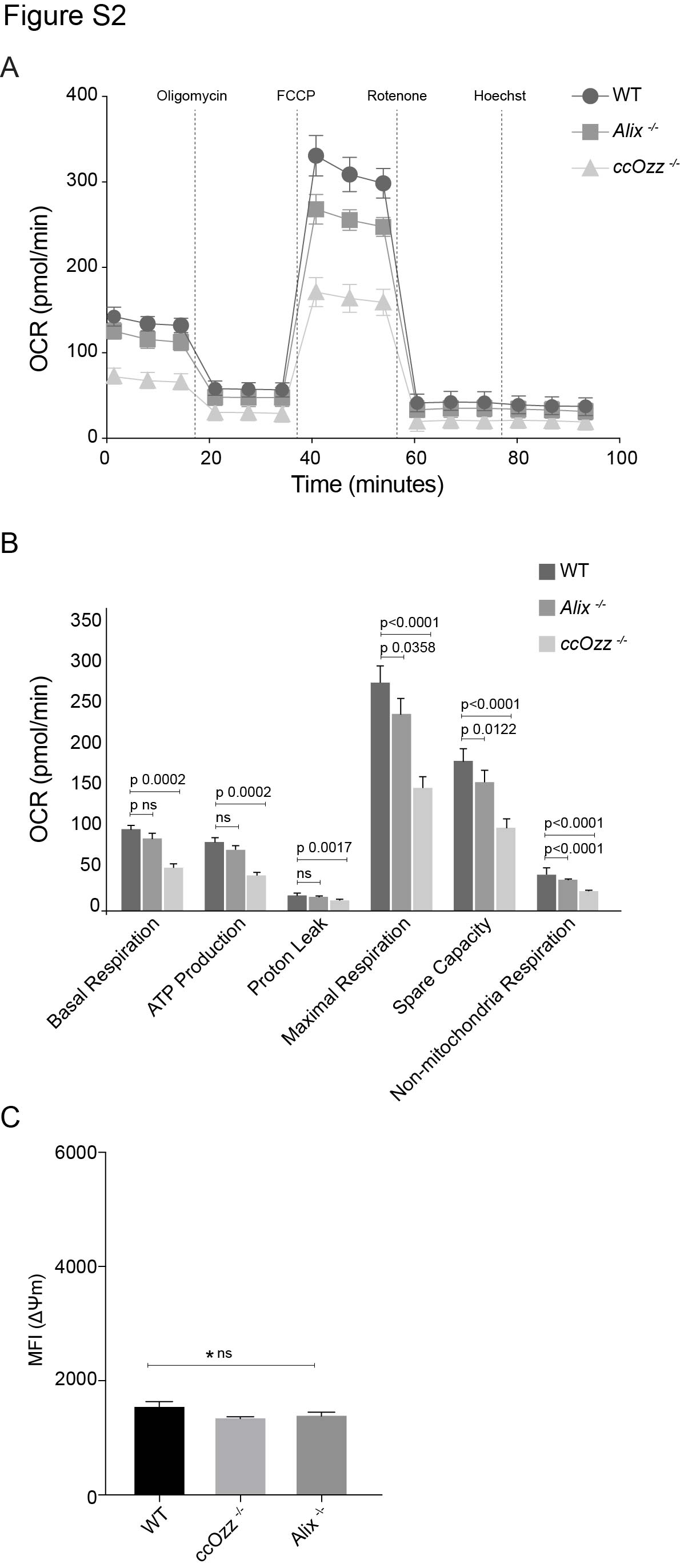

### Supplemental Figure 3

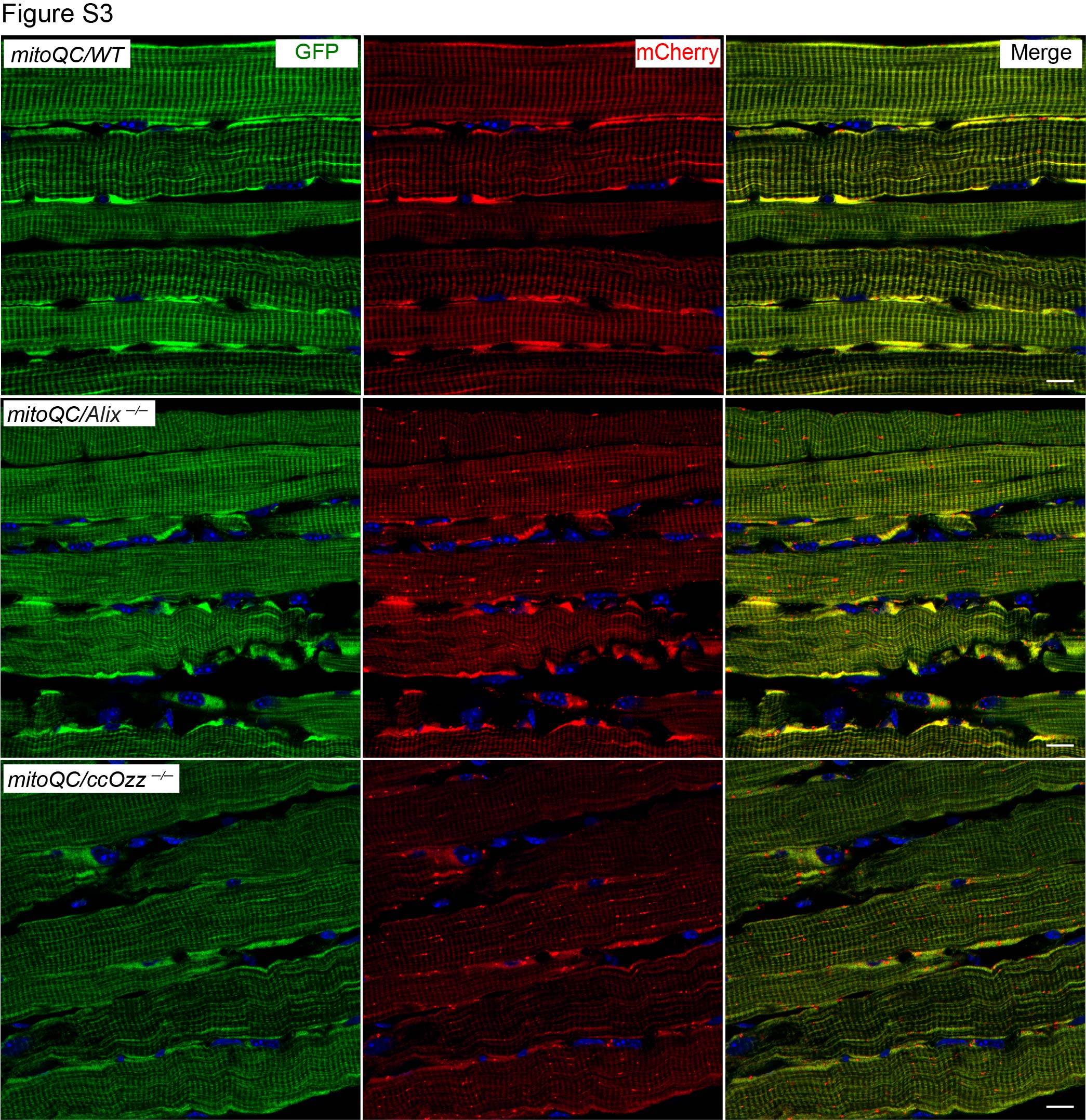

### Supplemental Figure 4

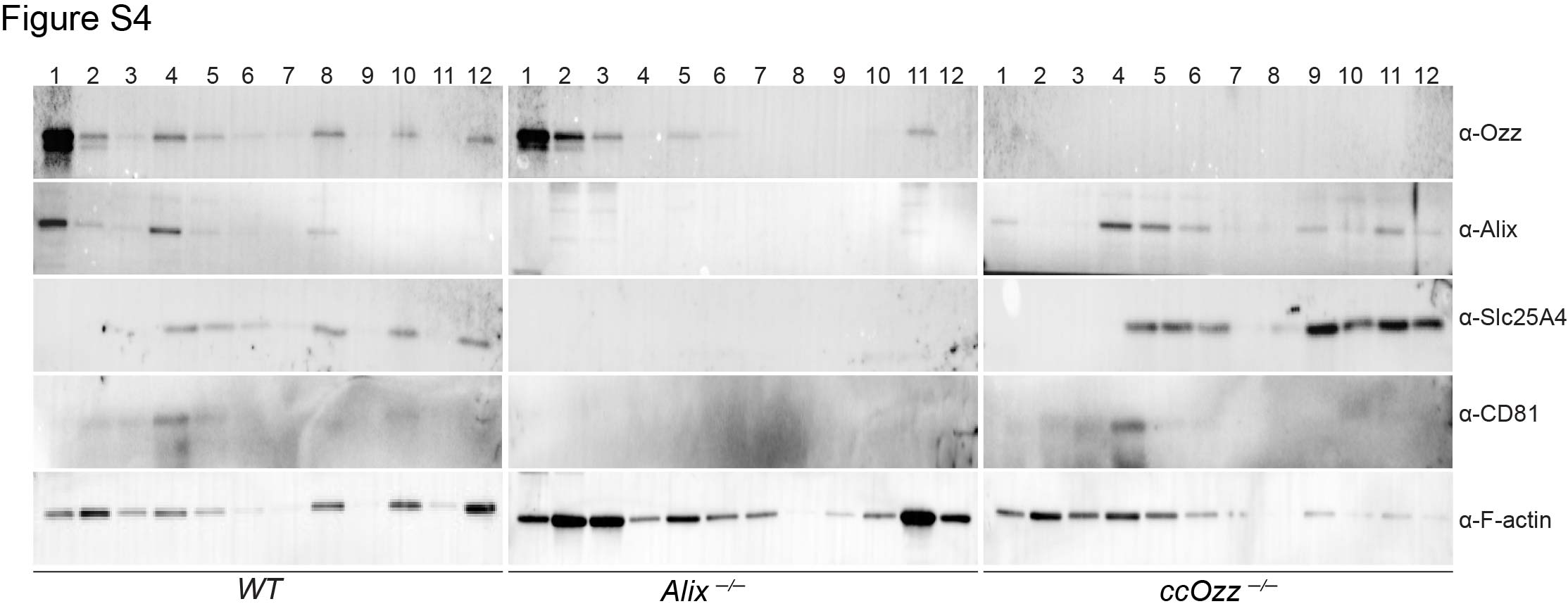
